# Bacteria rewire fungal antimicrobial gene expression in microbial arms races

**DOI:** 10.64898/2026.05.07.723608

**Authors:** Jinyi Zhu, Fantin Mesny, Shan Chen, Yukiyo Sato, Anton Kraege, Wilko Punt, Mehrnaz Zaeifi, Saifei Liu, Kathrin Wippel, Stéphane Hacquard, Bart P. H. J. Thomma

## Abstract

Microbial communities are shaped by antagonistic interactions, yet how competitors evade antimicrobial strategies in a community context remains unclear. Fungi deploy antimicrobial proteins (AMPs) to suppress bacterial niche competitors, but how bacteria counteract these defences is unknown. Here we show that soil microbiota suppress fungal AMP expression, revealing a previously unrecognized mode of intermicrobial interference. In the soil-borne pathogen *Verticillium dahliae*, AMP genes are highly conserved yet broadly repressed in the presence of natural microbial communities. This repression extends across diverse fungi, suggesting a widespread phenomenon. Correlation analyses of AMP expression and community compositions linked the bacterial genera *Leptothrix* and *Devosia* to repression of the *V. dahliae* AMPs VdL1 and VdD1, respectively. Both AMPs antagonize these bacterial taxa, and microbiota-mediated repression of *VdL1* reduces its contribution to fungal soil colonization. Together, our findings reveal that bacteria can evade fungal antagonism by suppressing AMP deployment, highlighting community-level control of microbial competition.

## INTRODUCTION

All life on earth is fundamentally interconnected, as organisms continuously interact across environments. These interactions range from beneficial and commensal relationships to intense antagonism, and frequently involve trophic interactions and metabolic exchanges. Ultimately, these interactions play crucial roles in shaping evolutionary adaptation and ecological relationships^1,2^. Plants live in close association with diverse microorganisms, collectively known as the plant microbiota, which predominantly comprises bacteria, fungi, and protists^3^. Together with their host, these microbial communities form the plant holobiont, a complex ecological entity^4^, in which host and microbes form tightly interconnected biological networks that influence each other’s physiology and ecological performance^5,6^.

Plant pathogens secrete so-called “effectors”, which are typically small, secreted proteins that fulfil diverse functions to facilitate infection, including immune suppression^7^. In response, in a lineage-specific fashion, plants evolved to recognize effectors or their activities to re-install immune responses, often through intracellular nucleotide-binding leucine-rich repeat (NLR) receptors^8–10^. Such recognition exerts selection pressure on the pathogen to lose or modify recognized effectors, or otherwise perturb the re-installed immune responses, which poses selection pressure on the host again to evolve novel recognition specificities in turn. This perpetual dynamics with cycles of attack, interception, recognition and evasion is described as an evolutionary “arms race” between pathogens and plants, which can result in a high degree of co-evolution. This is reflected in typical gene-for-gene interactions that characterize many plant-microbe interactions^11–13^.

Under pathogen attack, plants also recruit protective microbes to limit disease progression, a phenomenon termed “cry for help”^14–17^. For example, cucumber plants infected by *Fusarium oxysporum* recruit beneficial *Bacillus amyloliquefaciens* to mitigate disease^18^. Such responses may leave lasting effects on soil microbiota, as beneficial microbes persist and contribute to disease-suppressive soils. Increasing evidence suggests that plants actively shape their microbiota through metabolic cues and immune signaling^3,19–22^, while microbiota members suppress disease through diverse mechanisms^3,23–25^. Accordingly, the plant microbiota constitutes a barrier that pathogens must overcome, recently termed the “extended immune system”^26^.

Various pathogens secrete antimicrobial proteins (AMPs) to modulate host-associated microbiota as a strategy to establish infection and promote host colonization^27–33^. For instance, the soil-borne fungal pathogen *Verticillium dahliae*, which causes vascular wilt diseases in more than 400 plant species, secretes several AMPs that suppress antagonistic microbes during host colonization to promote infection^27–29,33^. Moreover, the *V. dahliae* genome encodes hundreds of such AMPs, highlighting their importance in microbiota manipulation^34^. These AMPs are secreted not only during host colonization but also during life stages outside the host, including soil-dwelling stages^27,33^. However, AMP production is not restricted to plant pathogenic fungi, and many fungi are predicted to encode extensive AMP catalogs irrespective of lifestyle^34^. A significant proportion of these AMPs is conserved across fungi with various lifestyles, suggesting an ancient origin of these AMPs^34^. This broad distribution and conservation of AMPs suggest that AMP-mediated microbial antagonism constitutes a fundamental and longstanding feature of fungal biology.

Fungal AMPs often display selective antimicrobial activity, meaning that they inhibit only certain microbes^27–33^, thereby imposing selection pressure to evolve resistance strategies. Here, we assessed how bacterial members of soil microbiota deal with fungal-secreted AMP effectors and hypothesize that bacteria evolved to modulate fungal gene expression to protect themselves.

## RESULTS

### *Verticillium* AMPs are functionally diverse and highly conserved

The predicted *V. dahliae* secretome comprises 894 proteins, including 91 transmembrane proteins and 263 carbohydrate-active enzymes (CAZymes)^34^. Antimicrobial activity prediction using AMAPEC^34^ showed that 179 of 263 CAZymes (68.1%) possess antimicrobial potential and were classified as CAZyme-AMPs, whereas 84 lacked such activity (CAZyme-nonAMPs) (Fig. 1a). CAZyme-AMPs include glycoside hydrolases (GHs), carbohydrate esterases (CEs), polysaccharide lyases (PLs), and auxiliary activities (AAs), while CAZyme-nonAMPs span similar classes but additionally include glycosyltransferases (GTs) (Fig. 1b). Apart from the absence of GTs among CAZyme-AMPs, no significant enrichment or depletion of CAZyme classes was observed between the two groups (Fisher’s exact test, *P*>0.05).

**Fig. 1:**
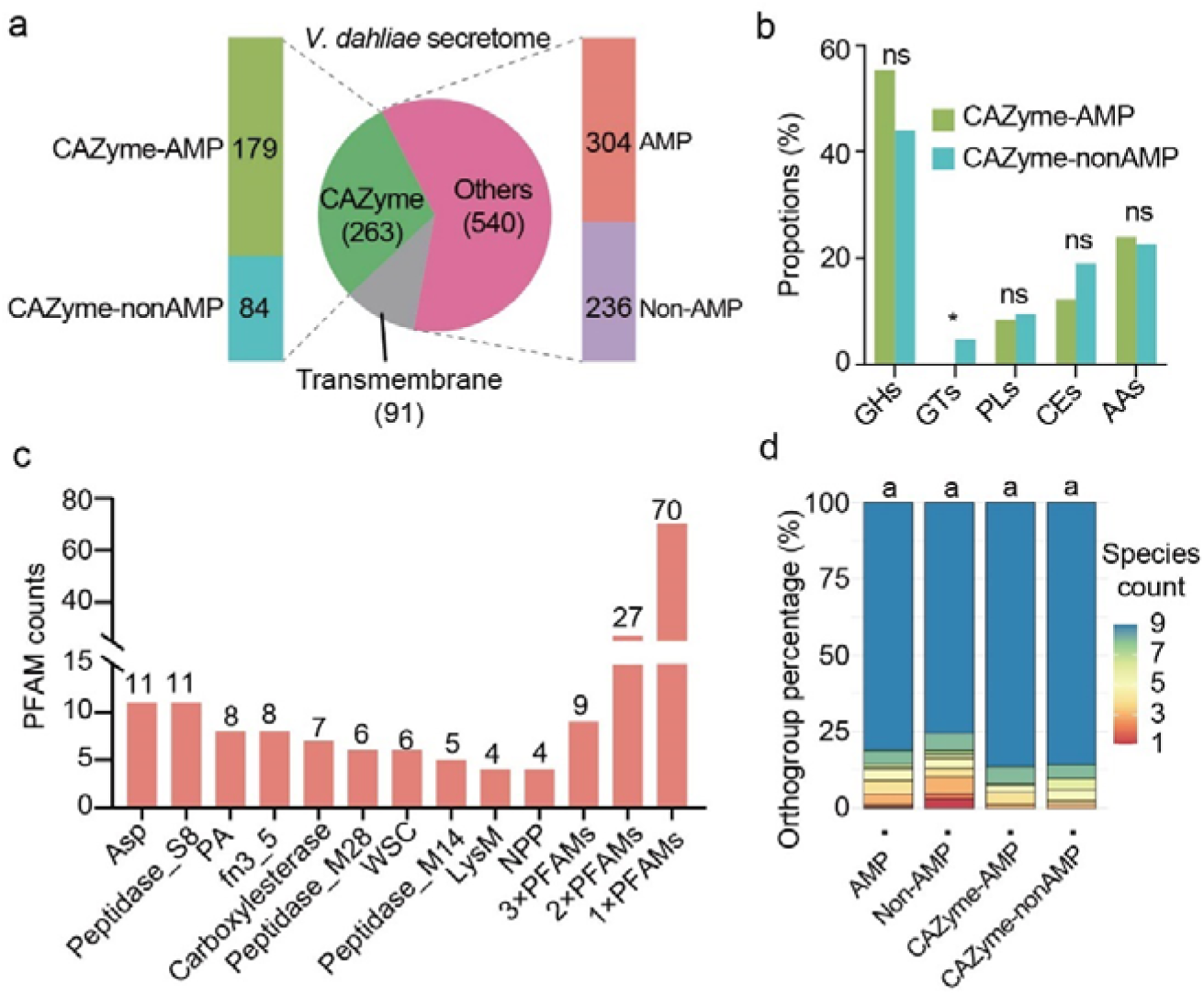
Characterisation of the *Verticillium dahliae* AMP gene catalog. **a,** Composition of the secretome of *V. dahliae* strain JR2 and antimicrobial activity prediction. The secretome was categorized into transmembrane proteins, carbohydrate-active enzymes (CAZymes), and other secreted proteins, with the number of proteins indicated for each category. Antimicrobial activity prediction was performed for CAZymes and other secreted proteins. **b,** CAZymes were classified into glycoside hydrolase (GH), glycosyltransferase (GT), polysaccharide lyase (PL), carbohydrate esterase (CE), and auxiliary activity (AA). Proportions (%) of each CAZyme class within the respective group are shown. Statistical differences were assessed using Fisher’s exact test (* p < 0.05; ns, not significant). **c,** Distribution of PFAM domains among 173 annotated *V. dahliae* AMPs. Numbers above the bars indicate the occurrence counts of each PFAM domain. The last three bars represent grouped PFAM domains that were detected three times (3× PFAMs), twice (2× PFAMs), or once (1× PFAMs). Abbreviations: Asp, aspartic protease domain; PA, protease-associated domain; fn3_5, fibronectin type III-like domain (family5); NPP, necrosis- and ethylene-inducing protein (NLP/NPP family); WSC, wall stress component; LysM, lysin motif. **d,** The degree of presence/absence variation of orthologs of AMP, non-AMP, CAZyme-AMP and CAZyme-nonAMP genes from *V. dahliae* strain JR2. Genes across nine *Verticillium* species were classified into orthogroups based on amino acid sequence conservation. Colors indicate the number of *Verticillium* species carrying at least one gene per orthogroup. Orthogroups were categorized according to the number of species in which they are present, and their relative proportions among all orthogroups are shown. Different letters indicate significant differences based on a Poisson generalized linear model followed by Tukey’s post-hoc test (two-sided p < 0.05) (n = 302 for AMP orthogroups, n = 234 for non-AMP orthogroups, n = 177 for CAZyme-AMP orthogroups, n = 84 for CAZyme-nonAMP orthogroups).

Besides transmembrane proteins and CAZymes, the *V. dahliae* secretome comprises 540 candidate effector proteins lacking clear functional annotation, including 304 predicted AMPs and 236 non-AMPs^34^ (Fig. 1a). Among the 304 AMPs, 173 contain recognizable PFAM domains, with 133 harboring a single domain and 40 containing two to four domains (Supplementary Table 1). Peptidase-related domains were most abundant, including Asp and Peptidase_S8 (each n=11), followed by PA (n=8), Peptidase_M28 (n=6), and Peptidase_M14 (n=5) (Fig. 1c). However, most PFAM domains were rare, with 70 occurring only once (Fig. 1c). Together, these results indicate that the predicted AMP set, including CAZymes and other types of effectors, comprises a highly diverse collection of proteins with divergent domain architectures and functional potentials.

To assess conservation of the secreted protein genes across the *Verticillium* genus^35^, we analyzed 42 high-quality genome assemblies, including 25 from *V. dahliae* and 17 from the remaining eight haploid species (Extended data Fig. 1). Genes were grouped into orthogroups based on sequence similarity, resulting 302 AMP, 234 non-AMP, 177 CAZyme-AMP, and 84 CAZyme-nonAMP orthogroups, most of which were present across all nine *Verticillium* species, with no significant differences in presence-absence patterns observed among the four categories (Fig. 1d).

### Soil microbiota repress fungal AMP gene expression

To test whether soil microbiota modulate fungal AMP gene expression, we used extracts from 10 previously characterized natural soils representing five major soil types (river clay, sea clay, sand, peat, and loam) that differ in physicochemical properties and microbiota composition^36^. Microbial growth in the extracts was confirmed by plating on tryptic soy agar (TSA; Extended data Fig. 2a, c), and 16S sequencing verified distinct bacterial community compositions (Extended data Fig. 2b). Microbe-free controls were generated by filter sterilization, which preserved soil-derived chemical composition while removing microorganisms (Extended data Fig. 2c).

Incubation of *V. dahliae* in soil extracts and filtrates resulted higher expression of three AMP genes (*VdAve1*, *VdAMP2*, and *VdAMP3*)^27,28^ in several filtrates than in the corresponding extracts, suggesting that their expression is suppressed by microbes present in these extracts (Fig. 2a, b). Intriguingly, the suppression pattern varied among soil extracts and AMP genes. For example, *VdAve1* was suppressed in peat 1 extracts, whereas *VdAMP2* and *VdAMP3* were not. Conversely, *VdAMP2* and *VdAMP3* were suppressed in extracts of river clays 1 and 2, whereas *VdAve1* was not. Expression of the housekeeping gene *VdTubulin* remained unchanged across treatments, excluding general transcriptional suppression (Fig. 2b). Together, these results suggest that soil microbiota can suppress *V. dahliae* AMP gene expression in a gene-dependent manner.

**Fig. 2:**
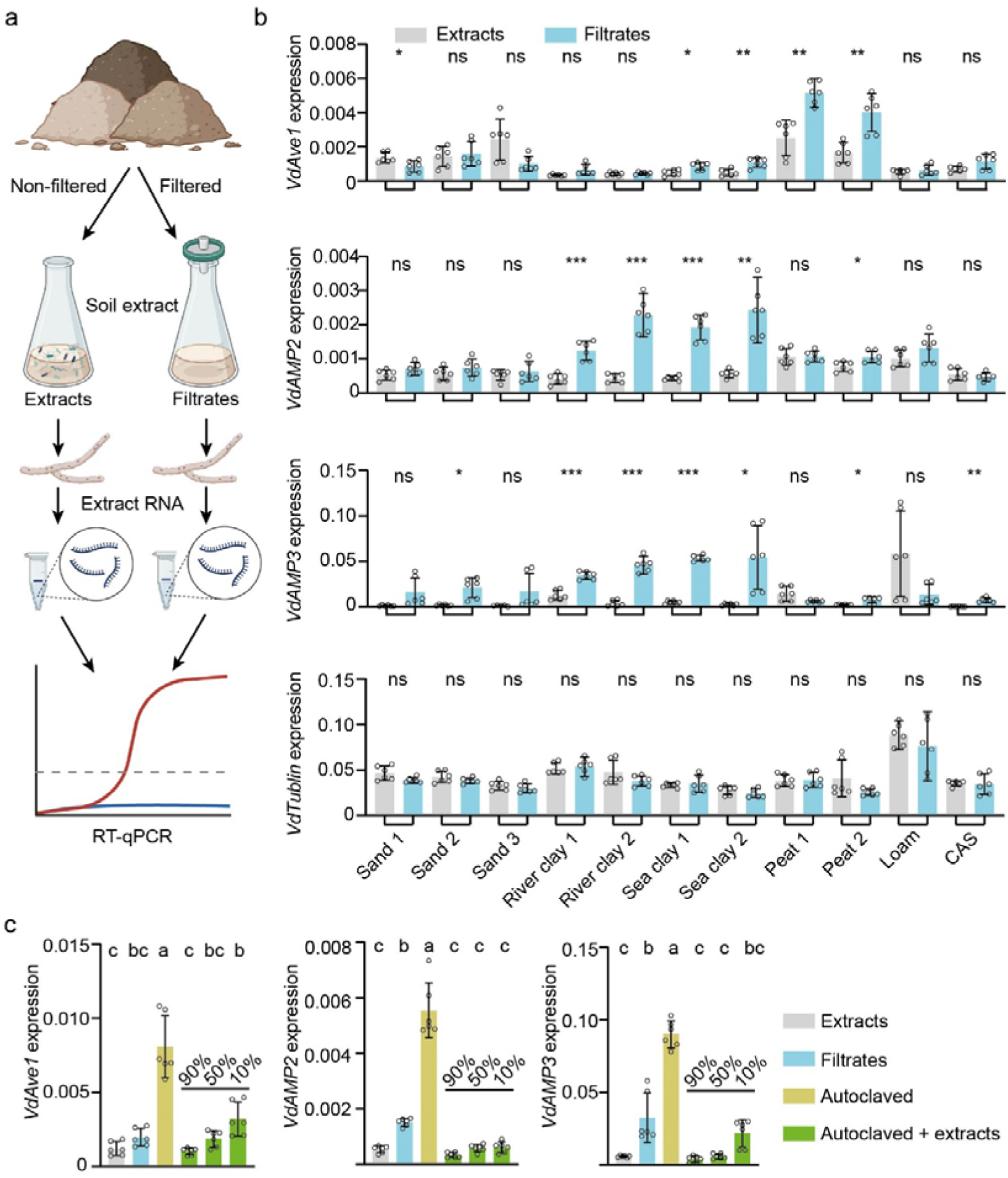
Soil microbiota suppress AMP gene expression in *Verticillium dahliae*. **a,** Schematic overview of the experimental design. *V. dahliae* mycelium was incubated for two days either in extracts from natural soil samples or sterile filtrates, followed by RNA extraction and RT-qPCR. **b,** Relative transcript levels of *VdAve1*, *VdAMP2*, and *VdAMP3* normalized to expression of the glyceraldehyde-3-phosphate dehydrogenase gene (*VdGAPDH*). *VdTubulin* (*VDAG_JR2_Chr3g08420*) expression was analyzed as negative control. Bars are shown with standard deviations (SD) and represent mean values of 6 independent biological replicates. Statistical significance was determined using Student’s t-test with Benjamini-Hochberg correction. (ns, not significant; * FDR < 0.05; ** FDR < 0.01; *** FDR < 0.001). **c,** Relative transcript levels of *VdAve1*, *VdAMP2*, and *VdAMP3* normalized to expression of the *VdGAPDH* and calculated using the 2^ΔCt method. Bars are shown with standard deviation (SD) and represent mean values of 6 independent biological replicates that are indicated by dots. Different letters represent significant differences (one-way ANOVA and Tukey’s *post hoc* test; P□<□0.05).

To further investigate the suppression of AMP expression, we autoclaved sea clay 2 extracts, in which all three AMP genes were suppressed (Fig. 1b), to eliminate living bacteria while retaining microbial components (Extended data Fig. 2d). Following incubation with *V. dahliae*, expression of all three AMP genes was strongly induced (Fig. 2c). Remarkably, addition of extracts to the autoclaved sample suppressed this induction (Fig. 2c), with only 10% extract sufficient to fully restore repression (Fig. 2c). These results suggest that microbial components can induce AMP gene expression, whereas specific soil microbes suppress fungal AMP gene expression.

To globally assess the impact of soil microbiota on *V. dahliae* gene expression, we performed RNA sequencing on mycelium incubated in soil extracts or corresponding filtrates. Principal component analysis (PCA) revealed that fungal transcriptomes differed substantially among extracts derived from different soils, irrespective of microbiota presence, indicating that soil-associated traits alone influenced fungal gene expression (PERMANOVA: R² = 71.21%, P < 10^−4^). Importantly, within each soil, clear separation between extract- and filtrate-treated samples demonstrated a pronounced effect of soil microbiota on fungal transcriptomes (PERMANOVA: R² = 6.62%, P < 10□□) (Extended data Fig. 3a). The number of differentially expressed genes varied markedly across soils (Extended data Fig. 3b), suggesting microbiota-specific transcriptional responses. To further characterize these responses, genes were classified into functional categories (housekeeping, primary metabolism, signaling and regulatory genes, AMP, CAZyme-AMPs and other secreted proteins), and the proportion of differentially expressed genes in each category was quantified. Housekeeping and primary metabolism genes, and to a lesser extent signaling and regulatory genes, showed limited differential regulation between extracts and filtrates treatments (Fig. 3a). Interestingly, AMP genes showed the most extensive differential regulation across soils and were significantly overrepresented among repressed genes in most soil extracts (Fig. 3a, b). Finally, as expected based on the differential expression of *VdAve1*, *VdAMP2* and *VdAMP3* (Fig. 2a, b), AMP genes exhibited differential expression responses to different soil microbiota (Fig. 3c).

**Fig. 3:**
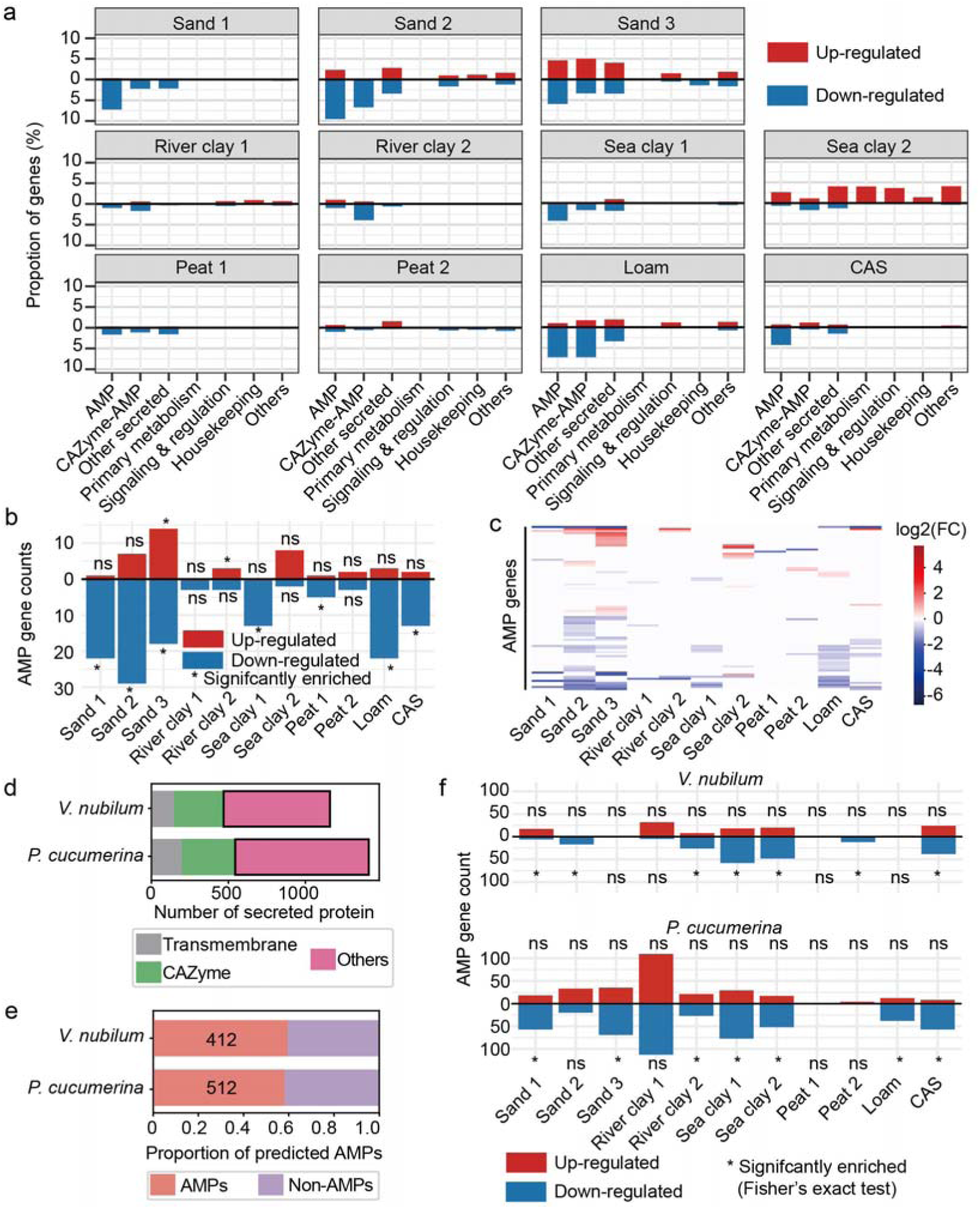
Soil microbiota repress AMP gene expression across fungi. **a,** For each soil sample, *V. dahliae* gene expression was compared between extracts and filtrates considering the functional categories antimicrobial protein (AMP), CAZyme-AMP, other secreted protein, primary metabolism, signaling and regulation, housekeeping, and other functions. Bars show the proportions of differentially regulated genes within each functional category, separated into up-regulated (red) and down-regulated (blue) genes, based on comparisons between non-filtered and filtered soil extracts for each soil sample. **b,** Numbers of differentially regulated AMP genes by the microbiota of each soil type. Asterisks indicate significant overrepresentation of AMP genes among differentially expressed genes in each soil, as assessed using Fisher’s exact test (ns, not significant). **c,** Heatmap of microbiota-regulated AMP gene regulation changes across soil samples. Log2 fold-changes were calculated for the extracts versus filtrates treatments per soil (DESeq2, adjusted *P*<0.05), with microbiota-induced up-regulation in red, and suppression in blue. **d,** Functional annotation of the predicted secretomes of *Verticillium nubilum* and *Plectosphaerella cucumerina*. Secreted proteins were classified as transmembrane proteins, CAZymes, proteins with other annotations, or proteins without annotation. The subsets of secreted proteins highlighted by black rectangles were used for antimicrobial effector prediction. **e,** Prediction of antimicrobial activity using AMAPEC, showing the proportions and numbers of predicted antimicrobial proteins (AMPs) within the filtered secretomes (excluding transmembrane proteins and CAZymes) of *P. cucumerina* and *V. nubilum*. **f,** Numbers of differentially regulated AMP genes by the microbiota of each soil type. Asterisks indicate significant overrepresentation of AMP genes among differentially expressed genes in each soil, as assessed using Fisher’s exact test (ns, not significant).

Given that fungi with diverse lifestyles produce AMPs^34^, we asked whether soil microbiota similarly repress AMP genes in other fungi. We therefore analyzed *Verticillium nubilum*, a saprophytic fungus^35^, and the soil-borne plant pathogen *Plectosphaerella cucumerina*^37,38^. We predicted the secretomes of both species and identified AMPs after excluding transmembrane proteins and CAZymes, resulting in 412 AMPs in *V. nubilum* and 512 in *P. cucumerina* (Fig. 3d, e). Interestingly, AMP genes were also repressed by soil microbiota in both species, with even larger numbers of AMP genes being repressed (Fig. 3b, f). In both fungi, AMP genes were significantly overrepresented among repressed genes (Fig. 4f), whereas other functional categories showed more limited changes in most soil extracts (Extended data Fig. 4a, b). Together, these results indicate that repression of fungal AMP genes by soil microbiota is a widespread and conserved phenomenon.

**Fig. 4:**
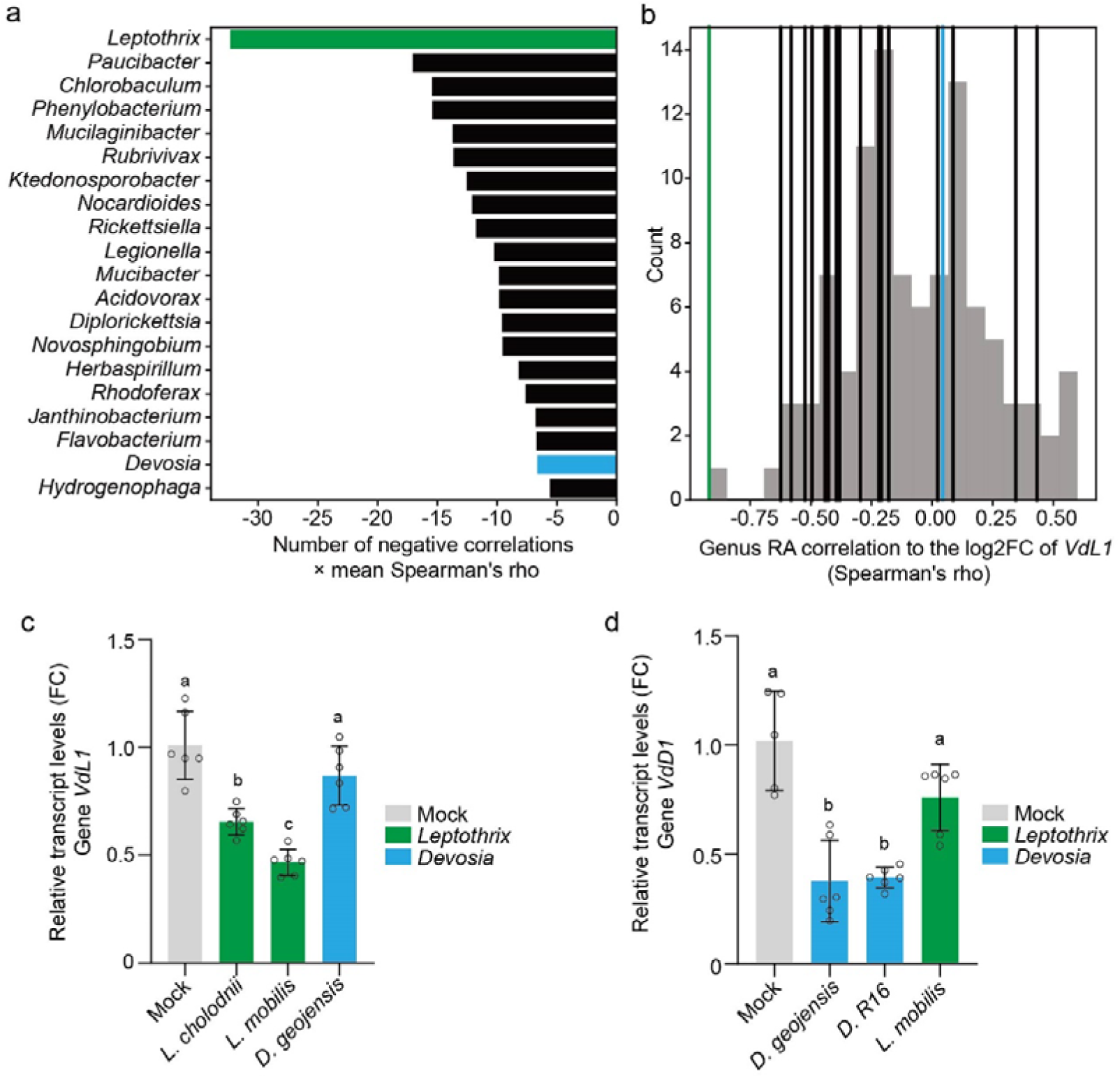
Specific bacteria suppress *Verticillium dahliae* AMP expression. **a,** top 20 bacterial genera are shown based on the index (number of negative correlations × mean Spearman’s rho). **b,** Distribution of Spearman’s correlation coefficients (Spearman’s rho) between the relative abundance of bacterial genus and the log2 fold change of *VdL1* expression. The x-axis shows correlation coefficients, and the y-axis indicates the number of bacterial genera within each correlation bin. Vertical lines indicate the positions of the 20 candidate genera identified in **a**. The green line marks *Leptothrix*, and the blue line marks *Devosia*. **c** and **d,** Relative transcript levels of AMP genes *VdL1* (**c**) and *VdD1* (**d**) in *V. dahliae* under different bacterial treatments. Gene expression levels were normalized to *VdGAPDH*. Bars represent mean values, and 6 individual dots indicate independent biological replicates. Different letters represent significant differences (one-way ANOVA and Tukey’s *post hoc* test; P□<□0.05).

### Specific bacteria suppress *V. dahliae* AMP expression

To evaluate potential associations between soil bacteria and fungal AMP gene repression, we performed correlation analyses linking bacterial taxon abundances with *V. dahliae* AMP gene expression across the 11 soil samples. To this end, relative abundances of bacterial genera were determined from 16S amplicon sequencing data (Supplementary Table S2), and genera exhibiting the highest variance in relative abundance across soils were ranked to retain the 100 most variable for analysis (Extended data Fig. 5a, b). *V. dahliae* AMP gene expression responses were quantified as log□ fold changes between extract and filtrate treatments for each soil, and correlated with the abundance profiles of the bacterial genera, yielding correlation matrices encompassing all pairwise relationships between 304 AMP genes and 100 genera (Extended data Fig. 5c). Interestingly, particular bacterial genera exhibited substantially more significant positive or negative correlations than others (Extended data Fig. 5d).

To identify candidate genera that may repress *V. dahliae* AMP expression, we calculated an index (number of significant negative correlations × mean correlation coefficient) and selected the top 20 negatively associated genera, with *Leptothrix* showing the strongest negative association (Fig. 4a). We tested whether *Leptothrix* represses four AMP genes with the strongest negative correlations (*VdL1*, *VdL2*, *VdL3*, and *VdL4*) (Extended data Fig. 6a). Indeed, both *Leptothrix cholodnii* and *Leptothrix mobilis* repressed *VdL1* expression, while *L. mobilis* additionally repressed *VdL2*, *VdL3*, and *VdL4* (Extended data Fig. 6b). To assess specificity of *VdL1* repression by *Leptothrix*, *Devosia* was used as a negative control, as it was predicted not to repress *VdL1* by the correlation analysis (coefficient approaching 0; Fig. 4b). Consistent with this, *Devosia* representatives did not repress *VdL1* expression (Fig. 4c) or *VdL2*-*VdL4* (Extended data Fig. 6b). However, *Devosia* abundance negatively correlated with a different set of AMP genes (*VdD1*, *VdD2*, *VdD3*, and *VdD4*), with *VdD1* showing the strongest association (Extended data Fig. 6c). Experimental validation confirmed that *Devosia* strains repressed *VdD1*, *VdD2*, and *VdD4*, but not *VdD3* (Extended data Fig. 6d). In contrast, *Leptothrix* did not repress *VdD1* (Fig. 4d), although it repressed *VdD2* and *VdD4* (Extended data Fig. 6d). Together, these results indicate that particular bacteria can suppress the expression of specific *V. dahliae* AMP genes.

### VdL1 and VdD1 orthologs occur widespread across fungi

As most *V. dahliae* AMP genes are widely present in the *Verticillium* genus (Fig. 1d), orthologs of *VdL1 and VdD1* were detected in all nine analyzed *Verticillium* species, including *V. nubilum* (Extended Data Fig. 7a), and were also identified in *P. cucumerina*. In *V. nubilum*, two *VdL1* orthologs (*VnL1-1* and *VnL1-2*) were identified, but neither showed negative correlation with *Leptothrix* abundance (Extended Data Fig. 7b, c). Similarly, *P. cucumerina* contains two *VdL1* orthologs, but neither is predicted to be secreted. For *VdD1*, three orthologs (*VnD1-1*, *VnD1-2*, and *VnD1-3*) were identified in *V. nubilum* and predicted as AMPs, of which *VnD1-1* and *VnD1-2* showed negative correlation with *Devosia* abundance (Extended Data Fig. 7d-f). In *P. cucumerina*, four *VdD1* orthologs (*PcD1-1*, PcD1-2, PcD1-3, and *PcD1-4*) were identified; *PcD1-3* is not predicted to be secreted and *PcD1-4* is not classified as an AMP. Neither *PcD1-1* nor *PcD1-2* showed negative correlation with *Devosia* abundance (Extended Data Fig. 7g, h).

To further assess the evolutionary conservation of *VdL1* and *VdD1*, we analyzed presence-absence of their orthologs across 150 fungal species spanning nine phylogenetic classes and eight lifestyle categories. *VdD1* orthologs were detected in all lineages, whereas *VdL1* orthologs were only absent from Glomeromycetes. Both were commonly detected across lifestyles, including pathogens and saprotrophs, although *VdL1* orthologs were not detected in arbuscular mycorrhizal fungi, endosymbionts, or orchid mycorrhizal fungi, and *VdD1* orthologs were not found in endosymbionts (Extended Data Fig. 7i).

### Antimicrobial activity of repressed AMPs

To gain insights into potential functions of VdL1 and VdD1, we examined their protein features. VdL1 harbors a predicted peptidase domain, whereas VdD1 lacks annotated PFAM domains (Supplementary Table S1). Structural analysis revealed that the C-terminal region of VdL1 shows similarity to a metallopeptidase M28 fold, whereas no structural similarity to known protein folds was detected for VdD1 (Extended data Fig. 7j, k).

Given that both proteins were predicted to have antimicrobial activity, VdL1 and VdD1 were heterologously expressed, and their antibacterial activities were tested against 12 phylogenetically diverse bacterial strains isolated from tomato plants^39^. Both exhibited selective antibacterial activity with distinct activity spectra (Fig. 5a; Extended data Fig. 8, 9). Based on the hypothesis that the repression of fungal AMP gene expression is a bacterial self-defense strategy, we tested whether VdL1 and VdD1 inhibit the growth of *Leptothrix* and *Devosia*. Interestingly, VdL1 inhibited both *Leptothrix* species, whereas VdD1 showed no detectable effect (Fig. 5b). Conversely, VdD1 inhibited *Devosia* strain R16, while VdL1 did not. Both proteins inhibited *D. geojensis* (Fig. 5c).

**Fig. 5:**
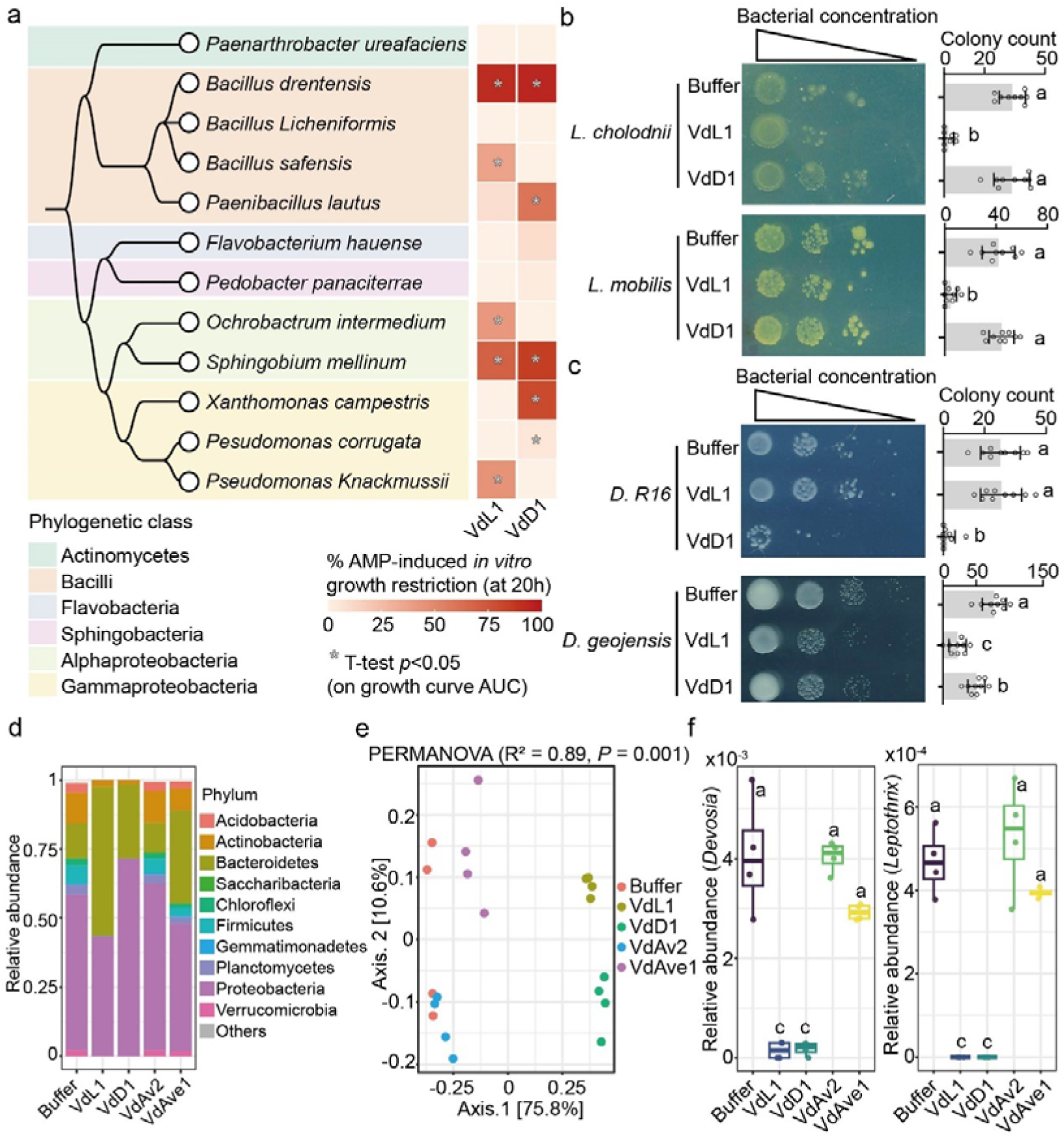
*Verticillium dahliae* effectors VdL1 and VdD1 exhibit selective antibacterial activity *in vitro* and reshape soil bacterial communities. **a,** Species phylogeny of a set of 12 bacteria generated with Taxallnomy^82^ and bacterial growth restriction induced by the presence of 8 µM of VdL1 or VdD1 calculated after 20 hours of incubation. Asterisks indicate significant differences compared with the buffer control as determined by a t test on growth-curve AUC (P < 0.05). Also see Extended Fig. 8-9. **b-c,** Spot dilution assays testing the antibacterial activity of VdL1 and VdD1 against *Leptothrix* (*L. cholodnii* and *L. mobilis*) and *Devosia* (strain R16 and *D. geojensis*). Bacterial suspensions were incubated with buffer, 8 µM VdL1 or VdD1 for 2 days and subsequently plated as a dilution series with bacterial concentrations decreasing from left to right. Colony counts are shown on the right. Data represent mean ± SD (n = 9). Different letters represent significant differences (one-way ANOVA and Tukey’s *post hoc* test; P□<□0.05). **d,** Relative abundance of bacterial communities at the phylum level in soil extracts after treatment with 1 µM of the indicated proteins (VdL1, VdD1, VdAv2, and VdAve1). **e,** Principal coordinates analysis (PCoA) based on Bray-Curtis dissimilarity of bacterial community composition in soil extracts following treatment with 1 µM of the indicated proteins. **f,** Relative abundance of *Devosia* (left) and *Leptothrix* (right) in soil extracts after treatment with 1 µM of the indicated proteins. For box plots (**d** and **f**), the center line indicates the median. Four individual data points represent biological replicates. Different letters represent significant differences (one-way ANOVA and Tukey’s *post hoc* test; P□<□0.05).

To assess their effects on microbial communities, purified VdL1 or VdD1 were added to soil extracts and compared with previously the characterized AMPs VdAve1 and VdAv2^27,33^. 16S amplicon sequencing revealed that AMP treatment altered bacterial community composition in soil extracts (Fig. 5d). Principal coordinate analysis (PCoA) showed clear separation between treatments, indicating distinct effects of different AMPs on microbial community structure (PERMANOVA: R² = 0.89, *P* = 0.001) (Fig. 5e). Interestingly, both VdL1 and VdD1 strongly reduced *Leptothrix* and *Devosia* relative abundances, whereas VdAve1 and VdAv2 had no significant effect (Fig. 5f). Together, these results demonstrate that VdL1 and VdD1 exhibit selective antibacterial activity.

### VdL1 contributes to *V. dahliae* soil colonization

Given that *VdL1* and *VdD1* exhibit selective antibacterial activity, we examined whether they contribute to fungal colonization in soil. A *VdL1* deletion mutant (*dVdL1*) and two independent complementation strains (*cVdL1-1* and *cVdL1-2*) were generated in *V. dahliae* strain JR2. All strains showed comparable growth on PDA (Fig. 6a, b).

**Fig. 6:**
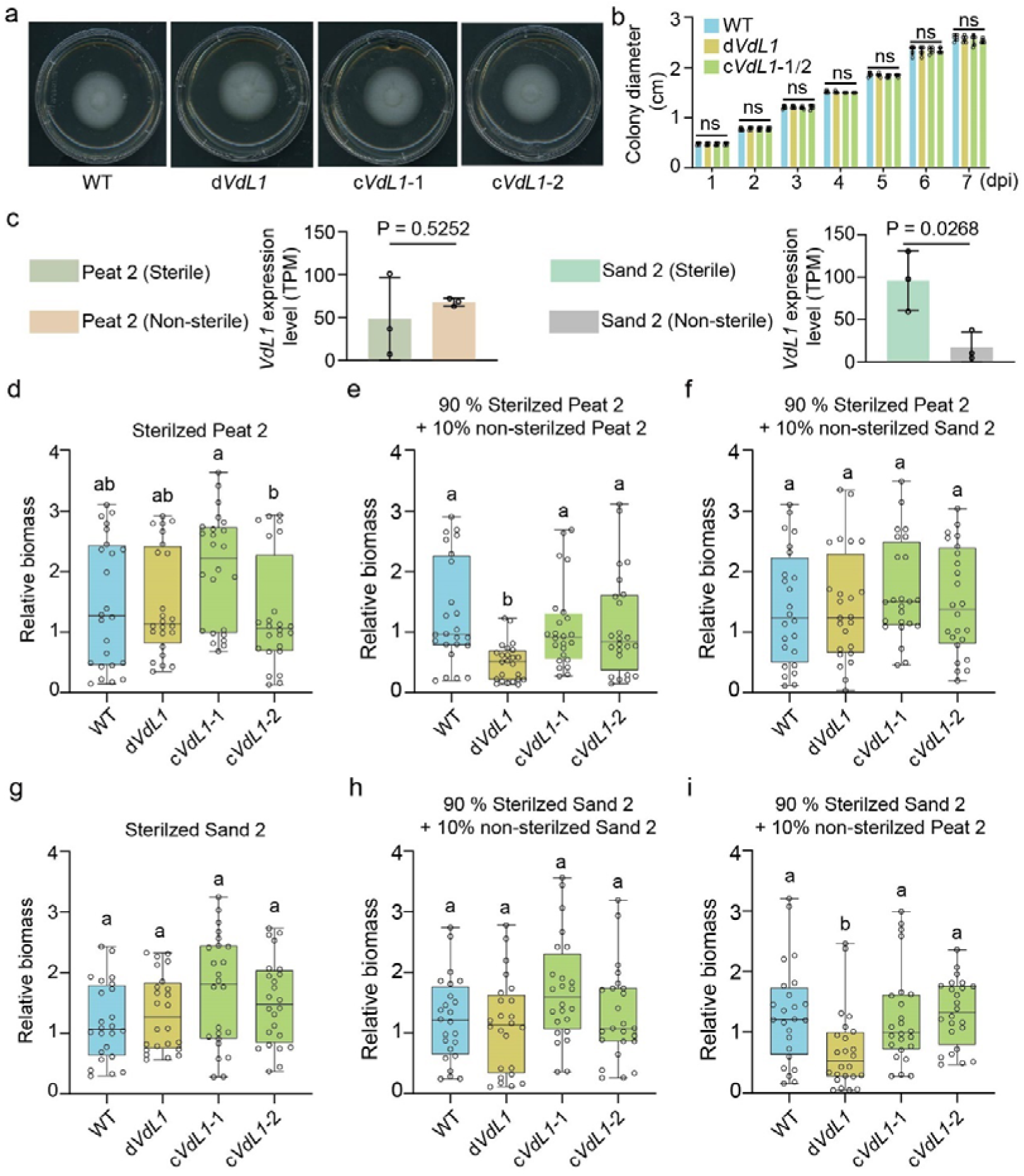
VdL1 contributes to *V. dahliae* soil colonization. **a**, Colony size of *V. dahliae* wild-type (WT), *VdL1* deletion (*dVdL1*), and complementation (c*VdL1*-1 and c*VdL1*-2) strains on PDA after 7 days. **b**, Colony diameter of WT, *dVdL1*, and complementation (c*VdL1*-1 and c*VdL1*-2) strains on PDA from day 1 to day 7 after inoculation. No significant differences were detected (one-way ANOVA with Tukey’s post hoc test). **c**, Expression of *VdL1* in Peat 2 and Sand 2 soils under sterile and non-sterile conditions based on RNA-seq data. Statistical significance was determined by Student’s *t*-test, with P values indicated in the figure. **d-i**, Relative biomass of WT, *dVdL1*, and complementation (c*VdL1*-1 and c*VdL1*-2) strains under the soil conditions indicated above each panel. Box plots show median (center line) and distribution; points represent biological replicates. Different letters indicate significant differences (one-way ANOVA with Tukey’s post hoc test, *P* < 0.05).

We selected Peat 2 soil, in which *VdL1* expression was not repressed, and Sand 2 soil, in which *VdL1* expression was significantly repressed (Fig. 6c). Each soil was autoclaved to generate sterile substrate, and subsequently added 10% non-autoclaved to reintroduce microbial communities and have recolonized soil. In sterile soil, no differences in colonization were observed among strains due to the absence of soil microbiota (Fig. 6d, g). Interestingly, in recolonized Peat 2 soil, the d*VdL1* mutant exhibited significantly reduced colonization compared to wild-type and complementation strains, indicating that VdL1 successfully targets antagonistic microbes that occur in Peat 2 soil (Fig. 6e). Intriguingly, no significant difference in colonization among strains was recorded in recolonized Sand 2 soil, which may be due to the occurrence of microbes that repress *VdL1* expression (Fig. 6h). Accordingly, the difference in soil colonization among strains in Peat 2 soil was not observed upon recolonization with Sand 2 microbes (Fig. 6f). Conversely, when sterile Sand 2 soil was recolonized with Peat 2 microbes, the d*VdL1* mutant showed significantly reduced colonization relative to the wild-type and complementation strains (Fig. 6i). Together, these results demonstrate that VdL1 contributes to *V. dahliae* soil colonization, and that this contribution is context-dependent as it can be repressed by soil microbiota.

## DISCUSSION

Intermicrobial interactions are pervasive in natural environments and play central roles in structuring microbial communities^40,41^. AMP-mediated antagonism represents a fundamental and ancestral feature of fungal biology, as fungi across lifestyles and phylogenies encode many evolutionarily conserved AMPs^34,42^. While the role of fungal AMPs in intermicrobial competition has been widely recognized, whether competing microbes actively defend against AMP-mediated antagonism has remained unclear. Our findings indicate that soil microbiota suppress fungal AMP gene expression across multiple fungi, and that specific bacterial taxa, exemplified by *Leptothrix* and *Devosia*, can repress AMP expression. Notably, the AMPs whose expression is suppressed by these bacteria display antibacterial activity against their repressors. Together, these observations suggest the existence of intermicrobial arms races, in which fungi deploy AMPs to inhibit bacteria, while bacteria counteract by repressing AMP expression.

Repressing antimicrobial expression represents one strategy by which bacteria avoid antagonism from other organisms, consistent with the diverse mechanisms that bacteria have evolved to counteract antimicrobial compounds depending on their mode of action^43,44^. These include modification of cell wall or cell membrane targets to prevent antimicrobial binding, secretion of proteases that degrade antimicrobial molecules, and active export of antimicrobials to reduce their concentration at the site of action^44–47^. Together, these strategies might allow bacteria to withstand antimicrobial pressure by interfering with antimicrobial activity. In contrast to these defense mechanisms, which operate at the level of antimicrobial action, our results proved that bacteria can repress fungal AMP gene expression as a defense strategy that may be particularly effective by reducing the production of multiple AMPs simultaneously, rather than counteracting each AMP individually. Consistently, most bacterial genera were correlated with the repression of multiple AMP genes (Extended data Fig. 5c, d). Accordingly, four AMP genes were confirmed to be repressed by *Leptothrix* (Extended data Fig. 6b). Importantly, *Devosia* repressed different AMP genes (Extended data Fig. 6d). Therefore, a bacterial community composed of diverse taxa may collectively repress a broader range of fungal AMPs than a single strain, suggesting that bacteria may compete collectively as a community with fungi.

Similar bacterial-induced transcriptional changes in fungi have also been reported. For example, antagonistic *Serratia* species induce transcriptomic reprogramming in the plant pathogenic fungus *Rhizoctonia solani* in an *in vitro* confrontation assay^48^. Likewise, *Ralstonia pickettii* downregulates fungal defense-related genes, including those involved in cell wall synthesis and reactive oxygen species (ROS) production, thereby shifting the interaction from antagonism toward commensalism^49^. In addition, antagonistic *Pseudomonas* and *Bacillus* strains trigger substantial transcriptional responses in the rice blast pathogen *Magnaporthe oryzae*^50^. However, these studies primarily focused on induced genes in fungi and did not assess whether secreted protein genes are preferentially repressed. In contrast, our study reveals that AMP genes represent the most strongly regulated functional category in *V. dahliae* by soil microbiota, while housekeeping and primary metabolism genes remain largely unchanged (Fig. 3a). Moreover, most differentially expressed AMP genes are repressed rather than induced (Fig. 3b). This repression-dominated pattern is also observed in *V. nubilum* and *P. cucumerina*, suggesting that bacterial-mediated repression of fungal AMP genes may represent a common feature of bacterial-fungal interactions. The underlying mechanisms remain unknown and warrant further investigation.

Bacteria also interact with phages and the mechanism of how bacteria defend bacteriophages infection have been well characterized^51^. Bacteriophages inject their genomes into bacterial cells, while bacteria deploy systems such as CRISPR-Cas to degrade invading DNA, and phages evolve countermeasures such as anti-CRISPR proteins. These interactions reflect ongoing intermicrobial arms races between bacteria and phages. Similarly, our findings suggest that bacteria engage in arms races with fungi. In many environments, including soil, bacteria and fungi compete for limited resources such as organic carbon, generating strong selective pressures that favor antimicrobial production by both partners^52^. Fungal AMPs represent potent weapons in this competition. A subset of fungal AMPs is deeply conserved across the fungal kingdom, suggesting that these proteins originated from ancient effector repertoires^32^. This evolutionary stability implies that AMPs provide a persistent fitness advantage, consistent with their selective activity against antagonistic organisms. For example, VdAve1 targets antagonistic members of the Sphingomonadales, whereas VdAv2 inhibits antagonistic *Pseudomonas* species^27,36^. However, such selective targeting is expected to impose strong survival pressure on affected bacteria. Our results indicate that some AMP-sensitive bacteria counteract this pressure by repressing fungal AMP expression (Fig. 4c, d). Notably, although many AMP genes are repressed in the presence of soil bacteria, a subset is simultaneously induced (Fig. 3b, f). While it remains unclear whether fungi actively upregulate alternative AMPs in response to bacterial repression, the induction may allow fungi to partially overcome bacterial counterstrategies.

Specific bacterial taxa can suppress fungal AMP gene expression, as exemplified by *Leptothrix* and *Devosia* (Fig. 4c, d). *Devosia* are known plant-beneficial bacteria that promote plant growth and suppress bacterial diseases^16^. Our findings suggest that *Devosia* may also influence plant-fungal interactions by modulating fungal AMP expression, potentially affecting fungal competitiveness in the plant-associated environment. Importantly, some fungal AMPs that are co-opted by plant fungal pathogens have dual roles in microbiota manipulation and host targeting^34^. Therefore, bacterial suppression of fungal AMPs represents an underexplored trait with potential for biological management of plant fungal diseases. Future work will be required to identify more such bacteria and to evaluate their effectiveness and stability under field conditions.

## MATERIALS AND METHODS

### Prediction of AMP genes

Antimicrobial effector catalogs were annotated in the genomes of *Verticillium dahliae* strain JR2^53,54^, *Verticillium nubilum* 397^55^, and *Plectosphaerella cucumerina* MPI-CAGE-AT-0016^56^ following a previously established procedure^34^. Secretomes were predicted from predicted protein sequences using SignalP v6.0^57^. Protein structures were predicted using ESM-Fold v1.0.3^58^, and candidate AMPs were identified using AMAPEC v1.0^34^, while excluding transmembrane proteins predicted with Tmbed v1.0.0^59^ and carbohydrate-active enzymes annotated with dbCAN v4.0^60^.

### *V. dahliae* AMP conservation analysis

To identify orthologs of AMP, non-AMP, CAZyme-AMP, and CAZyme-nonAMP genes of *V. dahliae* JR2 across *Verticillium* genomes, assembly qualities were assessed using BUSCO v5.8.3 with the eukaryota_odb12 dataset^61^. Genome assemblies were downloaded from NCBI using NCBI datasets v18.6.0^62^ and selected based on contig number (<300), complete BUSCO (>97%), multicopy BUSCO (<1%), and sequencing technology (Illumina, PacBio, or Nanopore with Illumina polishing).

Genes were annotated by homology using GeMoMa v1.9^63^ based on the *V. dahliae* JR2 annotation (VDAG_JR2v.4.0 in EnsemblFungi)^54^. *De novo* annotation was performed using BRAKER v3.0.8 pipeline C with fungal reference proteins from OrthoDB v11^64^. Annotations were merged using GffCompare v0.12.10^65^ with priority to homology-based predictions. Coding sequences were extracted using GeMoMa and translated with SeqKit v2.10.1^66^. Amino acid sequences were clustered into orthogroups using OrthoFinder v3.1.0^67^. Ortholog presence/absence variation was summarized using the R package tidyverse v2.0.0^68^.

Phylogenetic relationships of *Verticillium* species were inferred using REALPHY v1.13^69^. Genome sequences were fragmented into overlapping 50-bp reads and mapped to the *V. dahliae* JR2 reference genome using Bowtie2 v2.5.4^70^. Phylogenetic trees were constructed based on genome-wide SNPs using PhyML v3.3.20220408^71^ and visualized with ggtree v3.14.0^72^.

### Soil extract preparation

Soil samples were stored at 4 °C until use. Soil extracts were prepared by mixing 20 g soil with 100 mL sterile distilled water (1:5, w/v) and incubating at room temperature for 48 h. Soil particles were removed by centrifugation at 4,000 × g for 30 min, and supernatants were collected as soil extracts. Filtrates were obtained by passing extracts through 0.45 µm sterile filters. Autoclaved extracts were prepared by autoclaving soil extracts at 121 °C for 20 min. Mixtures of autoclaved and non-autoclaved extracts were prepared at ratios of 9:1, 1:1, and 1:9 (v/v).

### Determination of bacterial communities in soil extracts

Bacterial communities were analyzed by 16S rRNA gene amplicon sequencing. Ten mL of each soil extract was centrifuged at 12,000 × g for 30 min, pellets were resuspended in 50 µL sterile Milli-Q, transferred to PowerBead tubes, and DNA was extracted using the DNeasy PowerSoil Pro Kit (Qiagen, Hilden, Germany) following the manufacturer’s instructions (three biological replicates). The bacterial 16S rRNA gene was amplified using Phusion High-Fidelity DNA Polymerase (New England Biolabs, Ipswich, MA, USA) with primers 27F and 1139R. PCR reactions (25 µL) contained 0.5 µL polymerase, 5 µL HF buffer, 0.5 µL dNTPs, 1 µL of each primer, 1 µL template DNA, and nuclease-free water. PCR cycling consisted of 98°C for 2 min; 33 cycles of 98°C for 30 s, 56°C for 30 s, and 72°C for 45 s; followed by 72°C for 10 min. PCR products were purified using the Monarch Spin PCR & DNA Cleanup Kit (New England Biolabs, Ipswich, MA, USA). DNA concentration and quality were assessed by Qubit fluorometry and agarose gel electrophoresis. For each sample, 200 ng purified amplicon DNA was used for library preparation with the Nanopore sequencing kit SQK-NBD114.96 (Oxford Nanopore Technologies, Oxford, UK). Libraries were sequenced on an R10 flow cell using a MinION device for 72 h. Reads were processed and taxonomically classified using the EMU pipeline^73^.

### RNA preparation

*V. dahliae* JR2, *V. nubilum* 397, and *P. cucumerina* were cultured on PDA at 22°C for 7 days. Conidiospores (10□) were inoculated into 10 mL PDB and incubated at 22°C with shaking at 130 rpm for 2 days. Mycelia were collected with sterile Miracloth, washed twice with sterile Milli-Q, and transferred to flasks containing soil extract, filtrate, autoclaved extract, or their mixtures. Cultures were incubated for 2 additional days before RNA extraction using TRIzol reagent (Thermo Fisher Scientific, Waltham, MA, USA). RNA concentration and purity were measured using a NanoDrop spectrophotometer, and integrity verified by agarose gel electrophoresis. For sequencing, 10 µg RNA was treated with the TURBO DNA-free™ Kit (Thermo Fisher Scientific, Waltham, MA, USA). High-quality RNA (400 ng) was used for library preparation with the Optimal Dual-Mode mRNA Library Prep Kit and sequenced on a DNBSEQ-G400 platform (BGI, Shenzhen, China).

### RNA-seq analysis

Reads were quality filtered using Trimmomatic v0.39^74^ with parameters HEADCROP:10, TRAILING:20, AVGQUAL:20, and MINLEN:100. Filtered reads were mapped to fungal genomes using HISAT2 v2.2.1^75^. Genome assemblies and annotations of each fungus were used to build HISAT2 indexes. Alignments were sorted and indexed using samtools v1.18. Gene counts were obtained using featureCounts v2.0.6 with parameter -M^76^. Differential expression analysis was performed using DESeq2 v1.40.2^77^. Log2FoldChange values were shrunk using the apeglm method implemented in lfcShrink (apeglm v1.22.1)^78^.

### RT-qPCR

500 ng RNA was reverse-transcribed using HiScript® III RT SuperMix (Vazyme Biotech, Nanjing, China). cDNA was diluted ten-fold and qPCR performed using SsoAdvanced™ Universal SYBR® Green Supermix (Bio-Rad, Hercules, CA, USA) with gene-specific primers (Supplementary Table 3). Cycling conditions were 95°C for 3 min followed by 40 cycles of 95°C for 10 s and 60°C for 30 s. Melt curve analysis was performed from 65°C to 95°C. Relative expression was calculated using the ΔCt method with *VdGAPDH* as reference gene.

### Correlation of gene expression with bacterial abundances

Pairwise correlations were calculated between bacterial relative abundance (quantified by 16S metabarcoding sequencing) and AMP gene expression changes (log2FoldChange, presence vs absence of soil microbes) across 11 soil extracts. Spearman correlations between mean relative abundance and AMP gene log2FoldChange were calculated using stats.spearmanr() from the Python library Scipy v1.13.1^79^.

### Bacterial treatment to assess fungal AMP expression

*Leptothrix cholodnii* (BCCM: LMG 9467) and *Leptothrix mobilis* (DSMZ: DSM 10617) were cultured on M187 (1 g/L yeast extract, 1.5 g/L peptone, 0.2 g/L MgSO□·7H□O, 0.05 g/L CaCl□, 0.5 g/L ferric ammonium citrate, 0.05 g/L MnSO□·H□O, 0.01 g/L FeCl□·6H□O) agar. *Devosia geojensis* (DSMZ: DSM 19414) was cultured on TSB agar, and *Devosia R16*^80^ on TY (5 g/L tryptone, 3 g/L yeast extract, 0.87 g/L CaCl□·2H□O) agar. Bacterial cultures were grown in the corresponding media for 3 days at 25°C with shaking at 180 rpm. Cells were harvested by centrifugation (5000 × g, 3 min), washed twice with sterile Milli-Q, and resuspended in Sea clay 2 filtrate.

Conidiospores of *V. dahliae* JR2 were harvested from 7-day PDA cultures. A total of 10□ conidiospores were inoculated into 10 mL PDB and incubated at 22°C with shaking at 130 rpm for 2 days. Mycelia were collected using sterile Miracloth, washed twice with sterile Milli-Q, and transferred to flasks containing Sea clay 2 filtrate. Bacteria were added to a final OD_600_ of 0.05 and cultures incubated at 22°C with shaking at 130 rpm for 2 days before harvest.

### CRISPR–Cas9-mediated deletion and complementation of *VdL1* in *V. dahliae*

A CRISPR–Cas9-based genome editing protocol was used to delete *VdL1* in *V. dahliae* JR2, as previously described^34^. Two sgRNAs (Supplementary Table 3) targeting *VdL1* were designed using CRISPick (https://portals.broadinstitute.org/gppx/crispick/public). Donor DNA replacing *VdL1* with an mKate2 cassette flanked by 500 bp homology arms was assembled from PCR-amplified fragments of JR2 genomic DNA and *mKate2* using Phusion High-Fidelity DNA Polymerase (New England Biolabs, Ipswich, MA, USA). The fused fragment was cloned into pJET1.2 via the CloneJET PCR Cloning Kit (Thermo Fisher Scientific, Waltham, MA, USA), transformed into *E. coli* DH5α, and selected on LB agar with ampicillin. Correct inserts were confirmed by colony PCR and Sanger sequencing, and verified plasmids served as donor DNA for CRISPR–Cas9 editing. For complementation assays, two sgRNAs targeting *mKate2* were designed using CRISPick (Supplementary Table 3). The donor DNA consisted of the *VdL1* coding sequence flanked by 500 bp homology arms, which was amplified by PCR from *V. dahliae* JR2 genomic DNA and cloned into pJET1.2. Complementation transformation was subsequently performed in the *dVdL1* mutant background.

### Soil colonization assay

Peat 2 and Sand 2 soils were sterilized by double autoclaving. Soils were autoclaved once, mixed under sterile conditions, and autoclaved again. For recolonization treatments, non-autoclaved soil was mixed with autoclaved soil at a 1:9 ratio (w/w). Peat 2 or Sand 2 soils were recolonized with microbial communities from either soil type. Soil mixtures were incubated at room temperature in the dark for 2 days before fungal inoculation. For each treatment, 1 g soil was transferred to sterile 15 mL tubes.

Wild-type, VdL1 deletion (d*VdL1*) and VdL1 complementation (cVdL1-1, cVdL1-2) strains were cultured on PDA for 7 days. Conidiospores were harvested, washed twice with sterile Milli-Q, and 4 × 10□ conidiospores were inoculated into 30 mL PDB and cultured overnight with shaking at 60 rpm. Cultures were pelleted, washed twice with sterile Milli-Q, and resuspended in 4 mL sterile Milli-Q. 100 µL fungal suspension was added to each soil sample and incubated at room temperature in the dark. After 7 days, soils were homogenized and 250 mg subsamples used for DNA extraction with the DNeasy PowerSoil Pro Kit (Qiagen, Hilden, Germany). Spike-in plasmid (1 ng for each sample) was added to the CD1 lysis buffer for calibration^81^. DNA was eluted in 50 µL sterile Milli-Q.

Fungal biomass was quantified by qPCR using SsoAdvanced™ Universal SYBR® Green Supermix (Bio-Rad, Hercules, CA, USA). Cycling conditions were 95°C for 3 min followed by 40 cycles of 95°C for 10 s and 60°C for 30 s with fluorescence acquisition at each cycle. Melt curve analysis was performed from 65°C to 95°C with 0.5°C increments every 5 s. Relative biomass was calculated using the ΔCt method with the spike-in plasmid as reference.

### Protein production and purification

Coding sequences of *VdL1* and *VdD1* lacking signal peptides were amplified from *V. dahliae* JR2 cDNA using Phusion High-Fidelity DNA Polymerase (New England Biolabs, Ipswich, MA, USA). PCR fragments were cloned into the pET15b expression vector using the ClonExpress II One Step Cloning Kit (Vazyme Biotech, Nanjing, China). Constructs were transformed into *E. coli* DH5α, verified by colony PCR and Sanger sequencing, and transformed into *E. coli* BL21 for protein expression.

BL21 cultures were grown in 2×YT medium (16 g/L Bacto Tryptone, 10 g/L Bacto Yeast extract, 5 g/L NaCl) at 37°C until OD_600_ reached 1.5-2.0. Expression was induced with 1 mM IPTG followed by incubation at 42°C for 2.5 h. Cells were harvested and lysed in buffer containing 6 M guanidine-HCl, 10 mM Tris-HCl, and 10 mM β-mercaptoethanol (pH 8.0). Lysates were incubated overnight at 4°C and clarified by centrifugation (16,000 × g, 1 h). Proteins were purified by nickel affinity chromatography using His60 Ni Superflow Resin (Takara Bio, Shiga, Japan) on an ÄKTA go™ system (Cytiva, Marlborough, MA, USA). Protein fractions were identified by SDS-PAGE. Purified proteins were refolded by stepwise dialysis at 4°C (18 h per step, 1 L buffer) using the following buffers: Buffer 1: 4 M guanidine-HCl, 50 mM Tris-HCl, 0.5mM DTT, 10% glycerol, pH 8.0; Buffer 2: 3 M guanidine-HCl, 50 mM Tris-HCl, 0.5 mM DTT, 10% glycerol, pH 8.0; Buffer 3: 2 M guanidine-HCl, 50 mM Tris-HCl, 0.3 M L-arginine, 3 mM GSH, 0.3 mM GSSG, 10% glycerol, pH 8.0; Buffer 4: 1 M guanidine-HCl, 25 mM Tris-HCl, 0.3 M L-arginine, 20 mM NaCl, 2 mM GSH, 0.2 mM GSSG, 10% glycerol, pH 8.0; Buffer 5: 0.5 M guanidine-HCl, 25 mM Tris-HCl, 0.2–0.3 M L-arginine, 20 mM NaCl, 1 mM GSH, 0.1 mM GSSG, 10% glycerol, pH 8.0; Buffer 6: 10 mM potassium phosphate buffer, pH 7.6, 50 mM NaCl, 5% glycerol; Buffer 7: 10 mM potassium phosphate buffer, pH 7.4, 25 mM NaCl; Buffer 8: 10 mM potassium phosphate buffer. Final protein concentrations were determined using a Qubit 4 Fluorometer (Invitrogen, Waltham, MA, USA).

### *In vitro* microbial growth inhibition

Twelve tomato-associated bacterial strains^36^ were grown overnight in low-salt tryptic soy broth (ls-TSB) at 25°C with shaking at 180 rpm. Cultures were adjusted to OD_600_ = 0.05. 50 µL bacterial suspension was mixed with 50 µL purified protein (8 µM final concentration) or potassium phosphate buffer in 96-well plates. Plates were incubated at 25°C in a CLARIOstar® plate reader (BMG LABTECH, Germany) with double-orbital shaking every 15 min. OD600 was recorded every 15 min. For strain-specific assays, *L. cholodnii*, *L. mobilis*, *D. geojensis*, and *Devosia R16* cultures were adjusted to OD600 = 0.05. Bacterial suspensions (25 µL) were mixed with purified protein (8 µM) or buffer and incubated at 25°C with shaking for 2 days. Samples were serially diluted: *L. cholodnii* and *L. mobilis* (10×, 100×, 1,000×), *D. geojensis* (100×, 1,000×, 10,000×), and *Devosia R16* (10,000×, 100,000×, 1,000,000×). 5 µL of each dilution was spotted onto agar and incubated at 25°C for 5 days before imaging.

### AMP treatment of soil extract microbiota

Purified proteins were diluted to 2 µM in 10 mM potassium phosphate buffer. For each treatment, 25 µL Sea clay 1 extract was mixed with 25 µL protein solution or buffer control and incubated at 25°C with shaking at 180 rpm for 2 days. Total DNA was extracted using the DNeasy PowerSoil Pro Kit (Qiagen, Hilden, Germany). Four biological replicates were prepared for each treatment.

## ACKNOWLWDGEMENTS

F.M. acknowledges support from the Deutsche Forschungsgemeinschaft (DFG, German Research Foundation) through a Walter Benjamin Fellowship (Project ID: ME 6064/1-1; Project number: 508411006). Y.S. acknowledges funding from an Overseas Research Fellowship from the Japan Society for the Promotion of Science. S.C. and S.L. acknowledge funding from the China Scholarship Council. B.P.H.J.T. acknowledges funding from the Alexander von Humboldt Foundation in the framework of an Alexander von Humboldt Professorship funded by the German Federal Ministry of Education and Research, and further support from the DFG under Germany’s Excellence Strategy (EXC 2048/1; Project ID: 390686111) and from the DFG (Project ID: 458090666 / CRC1535/1). We thank Vittorio Tracanna for assistance with data management.

## AUTHOR CONTRIBUTIONS

JZ, FM, SH, and BPHJT conceived the project. JZ, FM, KW, SH and BPHJT designed the experiments. JZ, SC, AK, WP, MZ and SL performed the experiments. JZ, FM, YS, SC and BPHJT analysed the data. JZ and BPHJT wrote the manuscript. All authors read and approved the final manuscript.

