## Supplemental Figures for "Bacteria rewire fungal antimicrobial gene expression in microbial arms races"

**SUPPLEMENTARY FIGURES**

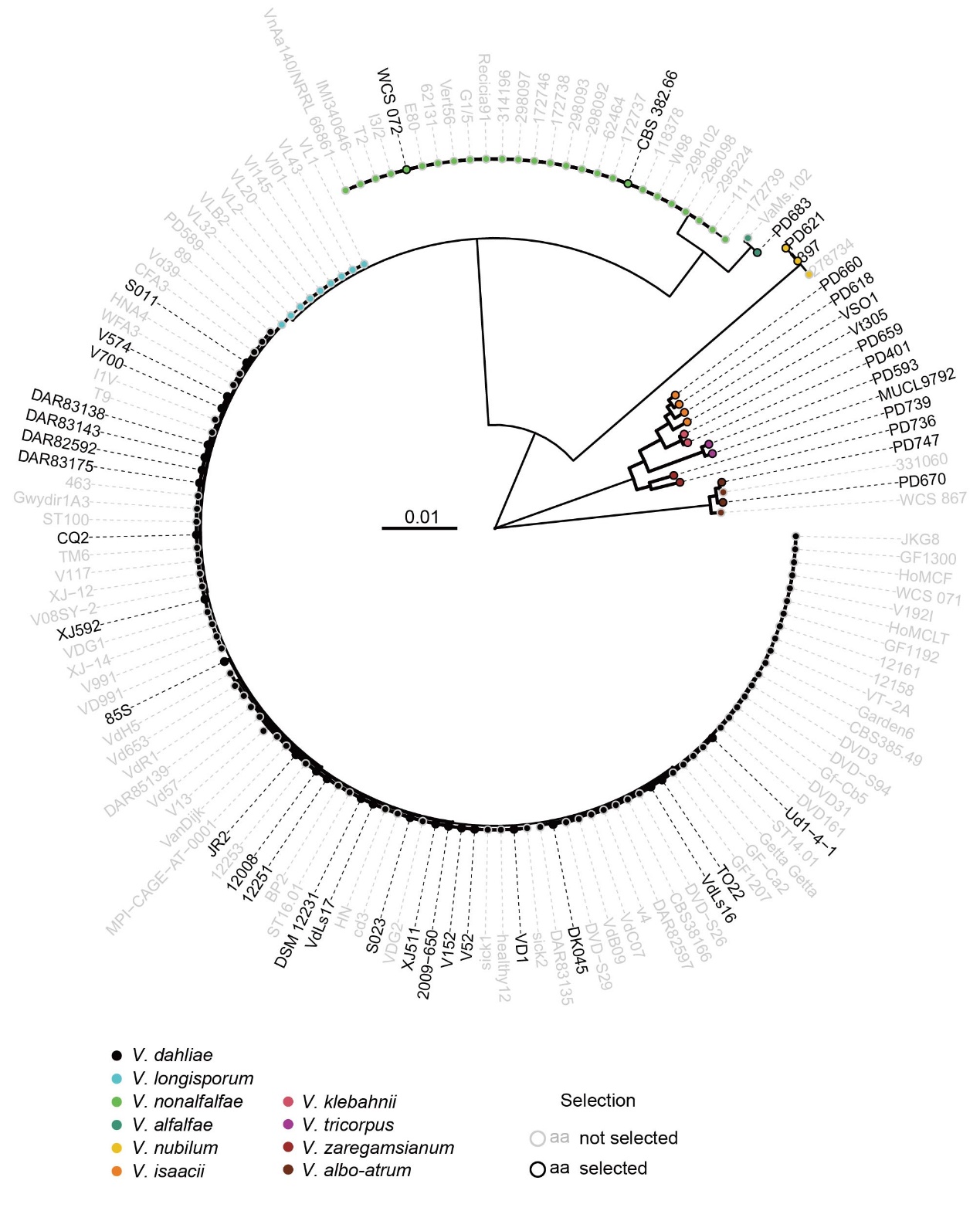

**Extended Data Fig. 1: Phylogenetic relationship of the ten *Verticillium* species.** The phylogenetic tree includes 143 strains (Supplementary Table 4) representing ten species within the genus *Verticillium* and was inferred from whole-genome sequence alignments. Tip colors indicate species identity, as shown in the legend. Strains highlighted in black were selected for downstream analyses shown in Fig. 1d.

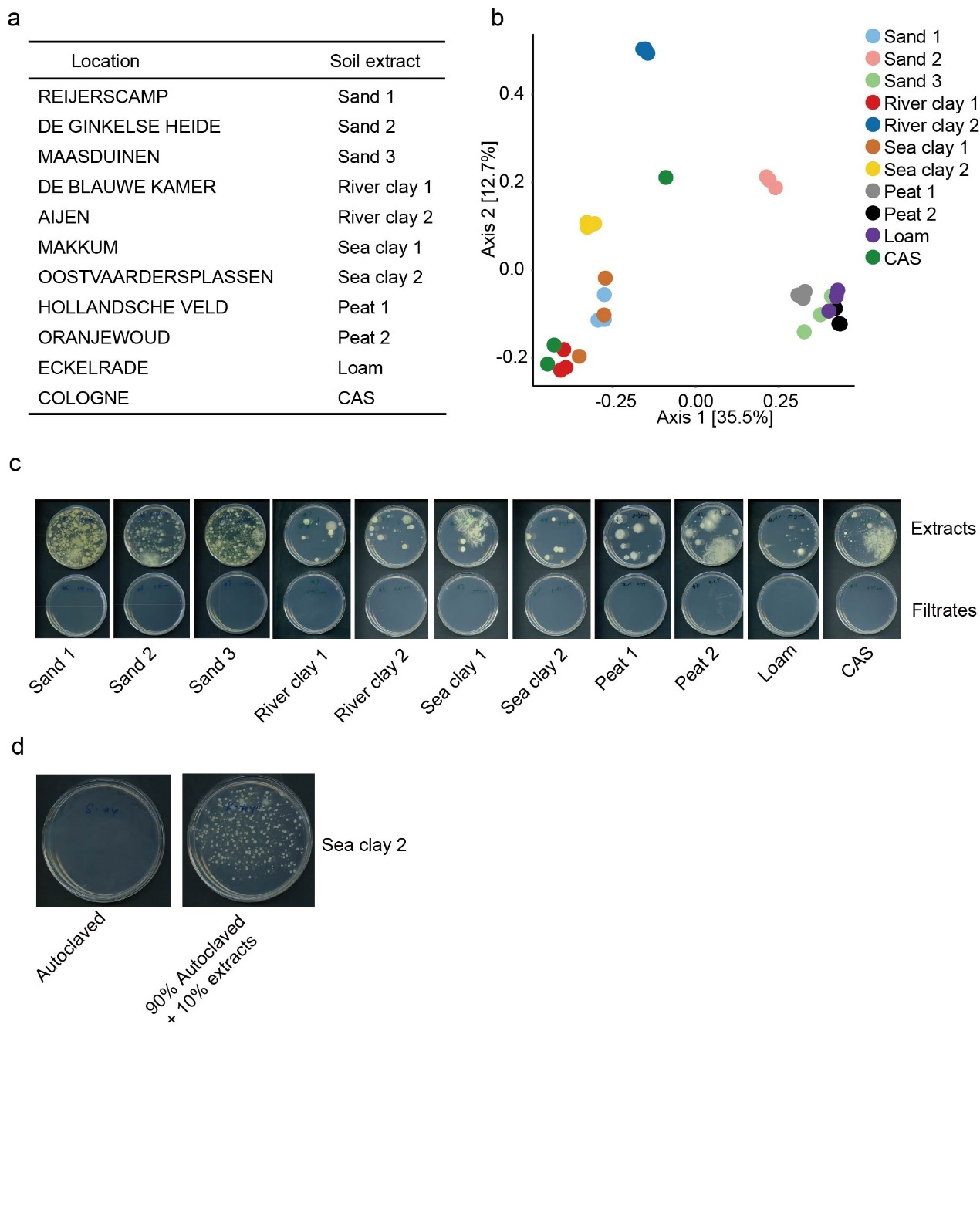

**Extended Data Fig. 2: Soil bacterial community compositions. a,** Overview of soil samples, including sampling location names and the corresponding soil extracts names. **b,** Principal coordinates analysis (PCoA) of soil extract bacterial community compositions based on 16S amplicon sequencing using Bray-Curtis dissimilarity. Three biological replicates are shown and the percentage of variance explained by each principal coordinate is indicated on the axes. **c,** Extracts and filtrates prepared from different soils were plated on TSB agar. **d,** Autoclaved Sea clay 2 extract and autoclaved soil extract supplemented with 10% extract were plated on TSB agar. Images were taken after 5 days of incubation at room temperature.

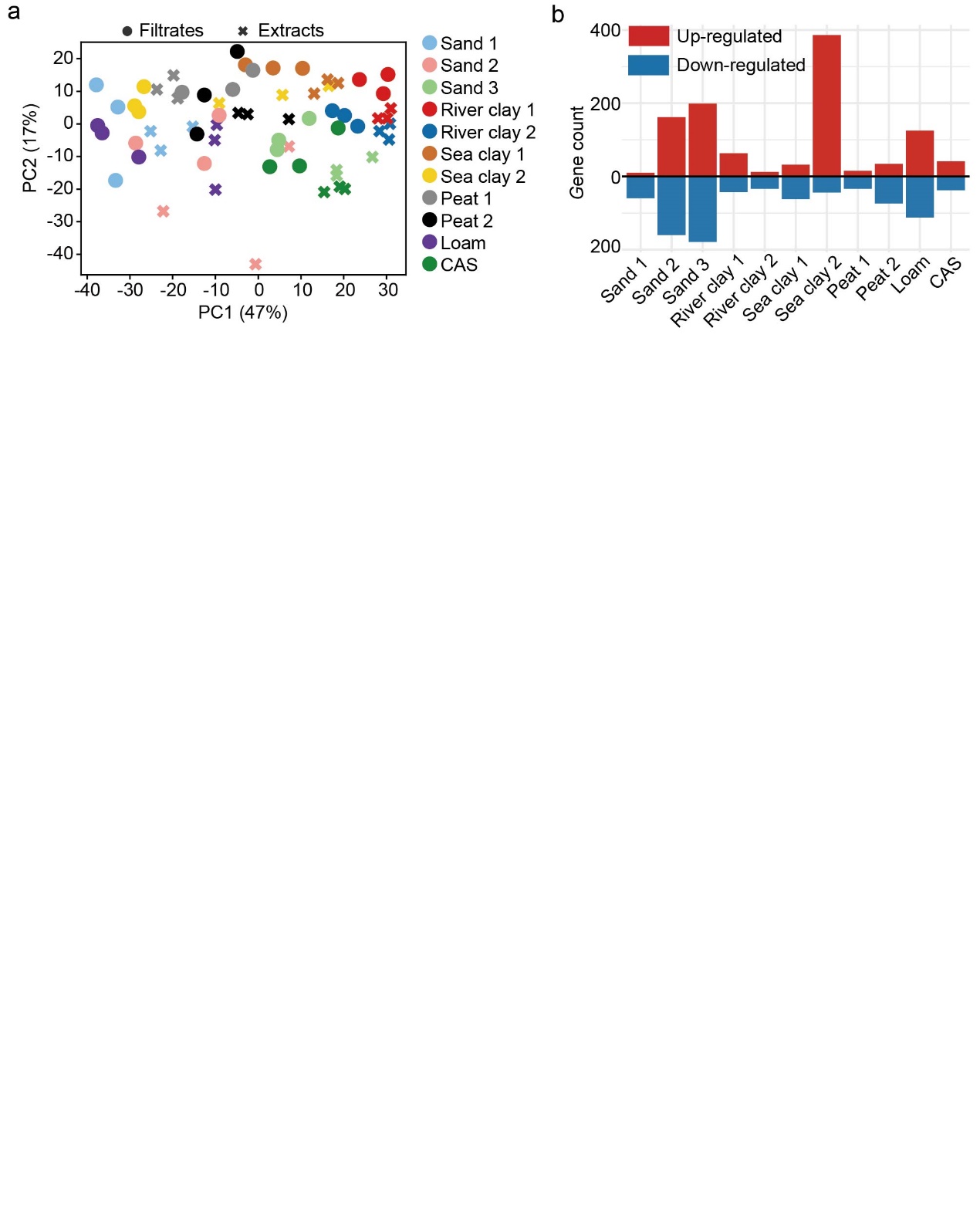

**Extended Data Fig. 3: Soil microbiota differentially affect *Verticillium dahliae* transcriptomes.** **a,** Principal component analysis (PCA) of *V. dahliae* transcriptomes upon incubation in soil extracts (crosses) or filtrates (circles). Each color represents a distinct soil sample. The percentage of variance explained by each principal coordinate is indicated on the axes. **b,** The number of differentially induced (red) or repressed (blue) genes in *V. dahliae* upon incubation in soil extracts when compared with filtrates (DESeq2, adjusted *P*<0.05).

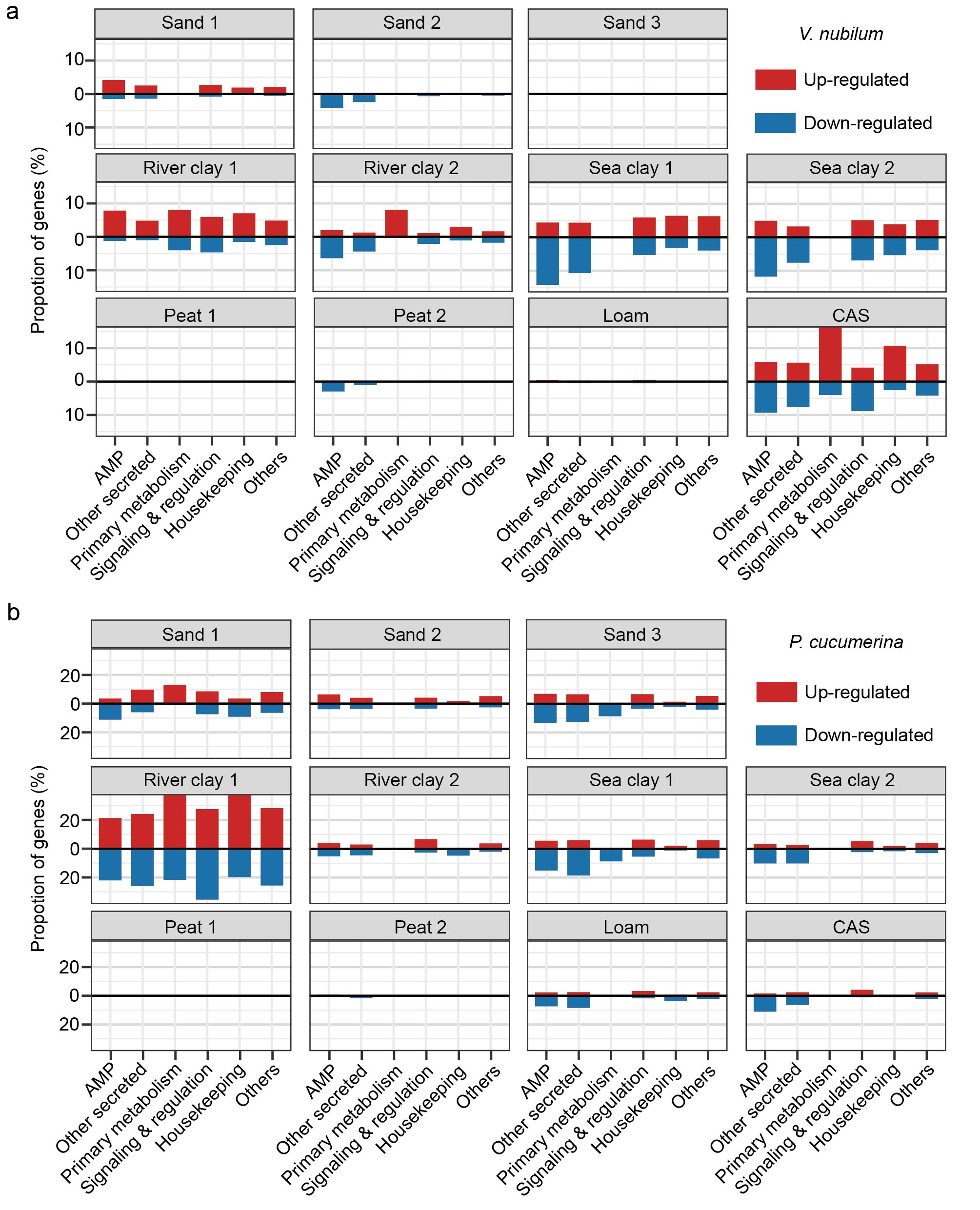

**Extended Data Fig. 4: Differentially expressed genes in *Verticillium nubilum* (a) and *Plectosphaerella cucumerina* (b).** Genes were grouped into functional categories: antimicrobial proteins (AMPs), other secreted proteins, primary metabolism, signaling and regulation, housekeeping, and other functions. Bars indicate the proportion of differentially expressed genes in each category, separated into upregulated (red) and downregulated (blue) genes, based on comparisons between soil extracts and filtrates for each soil sample.

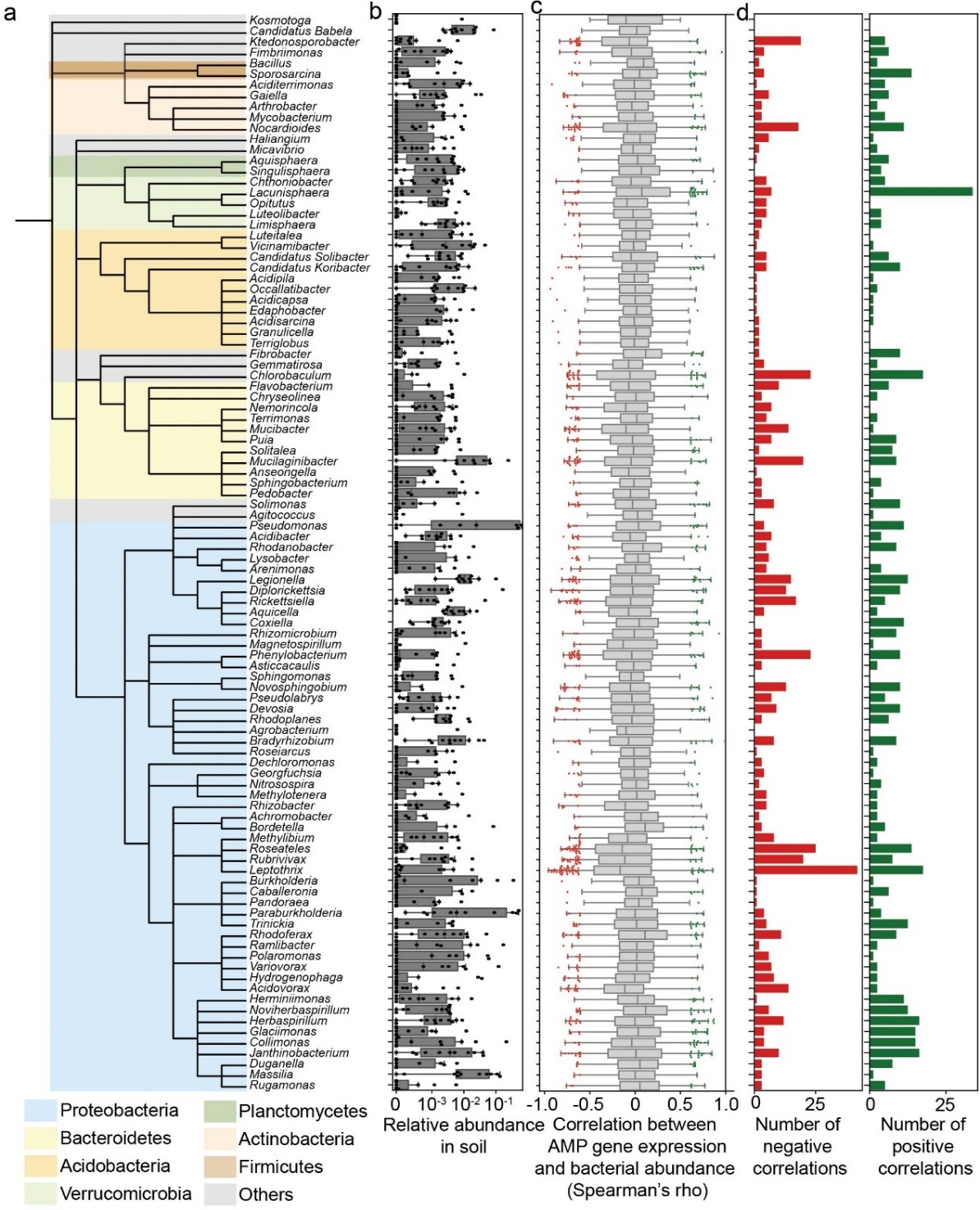

**Extended Data Fig. 5: *Verticillium dahliae* AMP gene repression correlates with bacterial abundance.** **a,** Phylogenetic tree of 100 bacterial genera included in the correlation analysis, with background colors indicating bacterial phyla. **b,** Relative abundance of the 100 most variable bacterial genera across soil extracts. **c,** Spearman’s rank test testing for correlation between AMP gene expression changes caused by soil microbiota (log2 fold change) and the relative abundances of bacterial genera across soils. Each dot represents a significant correlation (P<0.05). Red dots indicate negative correlations (higher bacterial abundance associated with lower AMP gene expression, rho<0), whereas green dots indicate positive correlations (higher bacterial abundance associated with higher AMP gene expression, rho>0). **d,** Number of negative (red bars) and positive (green bars) correlations between bacterial genera and AMP gene expression.

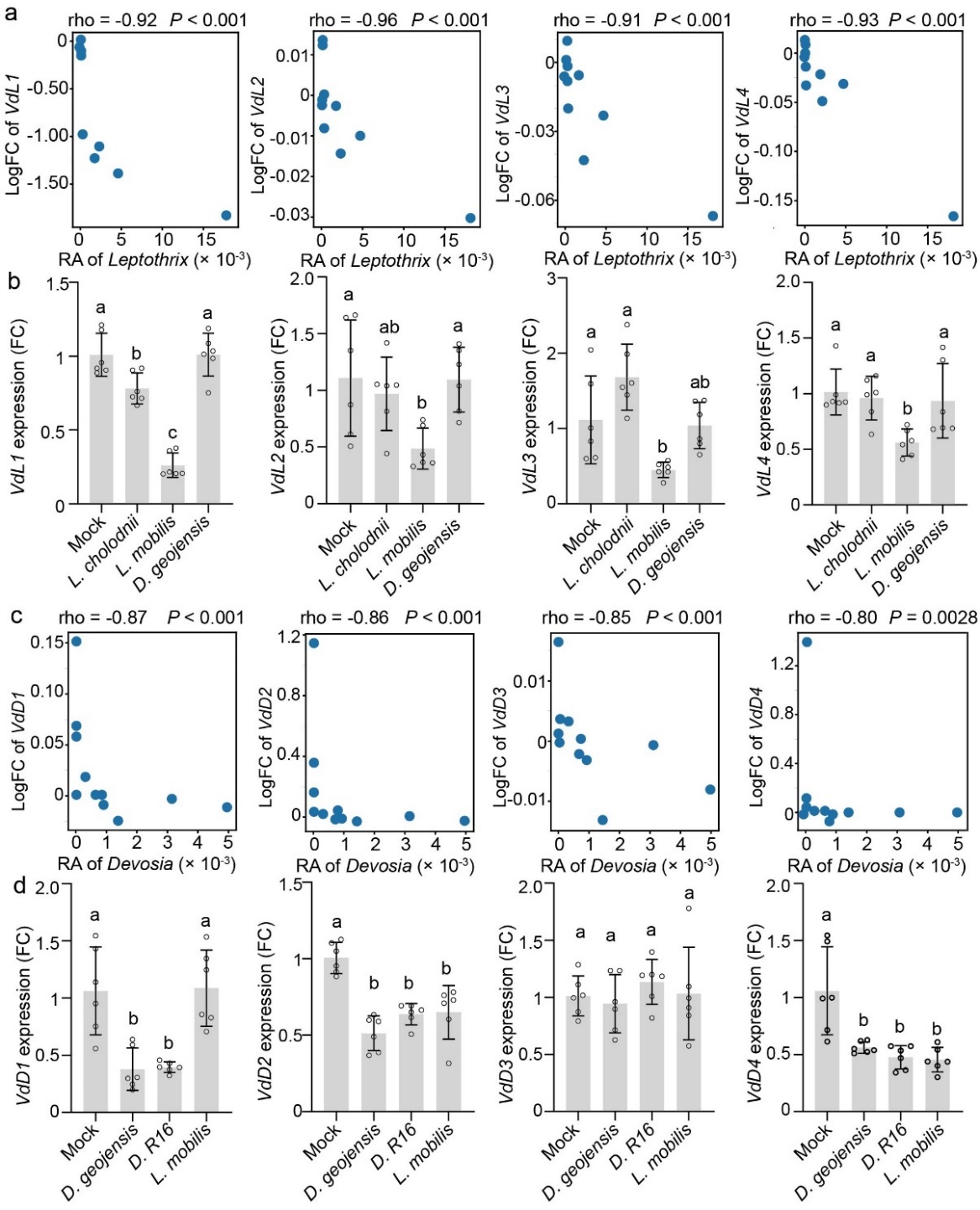

**Extended Data Fig. 6: AMP gene expression correlated with relative abundance of *Leptothrix* or *Devosia*. a,** Correlation (Spearman’s rank test) between the relative abundance of *Leptothrix* and the log₂ fold change (log₂FC) of the AMP genes *VdL1*, *VdL2*, *VdL3*, and *VdL4*. **b,** Relative transcript levels of the AMP genes *VdL1*, *VdL2*, *VdL3*, and *VdL4* in *V. dahliae* upon bacterial treatment. **c,** Correlation (Spearman’s rank test) between the relative abundance of *Devosia* and the log₂ fold change (log₂FC) of the AMP genes *VdD1*, *VdD2*, *VdD3*, and *VdD4*. **d,** Relative transcript levels of the AMP genes *VdD1*, *VdD2*, *VdD3*, and *VdD4* in *V. dahliae* upon bacterial treatment. Gene expression levels were normalized to *VdGAPDH*. Bars represent mean values, and dots indicate six independent biological replicates. Different letters represent significant differences (one-way ANOVA and Tukey's *post hoc* test; P < 0.05).

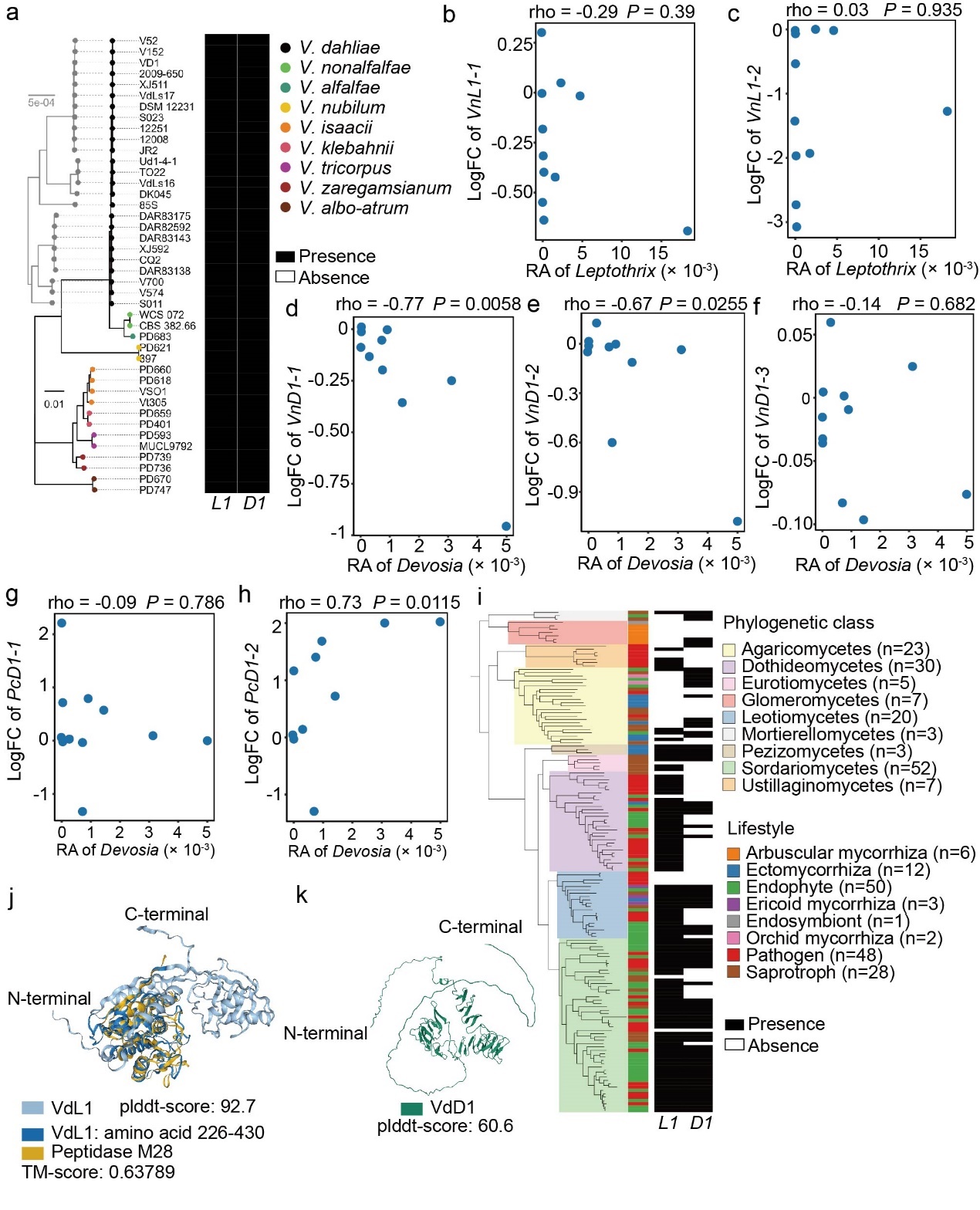

**Extended Data Fig. 7: VdL1 and VdD1 are ancient fungal effectors.** **a,** Presence/absence of L1 and D1 across the *Verticillium* genus. The phylogenetic tree shown in black represents relationships among 42 strains from nine *Verticillium* species inferred from whole-genome alignments, whereas the grey tree provides higher-resolution relationships among *V. dahliae* strains. **b-h,** Correlations (Spearman’s rank test) between bacterial relative abundance and AMP gene expression. Each panel shows the correlation between the relative abundance (RA) of the indicated bacterial genus (x-axis) and the log₂ fold change (log₂FC) of the indicated AMP gene (y-axis). Correlation coefficient (rho) and *P*-value are shown in each panel. **i,** Conservation of *L1* and *D1* across 150 fungal genomes based on orthology prediction. The phylogenetic tree illustrates the composition of the genomic dataset, with phylogenetic classes and fungal lifestyles annotated in colors. The presence or absence of orthologs of VdL1 and VdD1 is indicated by black and white squares, respectively. **j-k,** Predicted three-dimensional structures of VdL1 (**j**) and VdD1 (**k**) without signal peptides and superposition of the C-terminal region of VdL1 (amino acids 226–430) with a representative peptidase M28 structure, showing a peptidase M28-like fold. Structures are generated using AlphaFold, with overall predicted local distance difference test (plddt) scores indicated for each structural. The template modeling score (TM-score) is indicated.

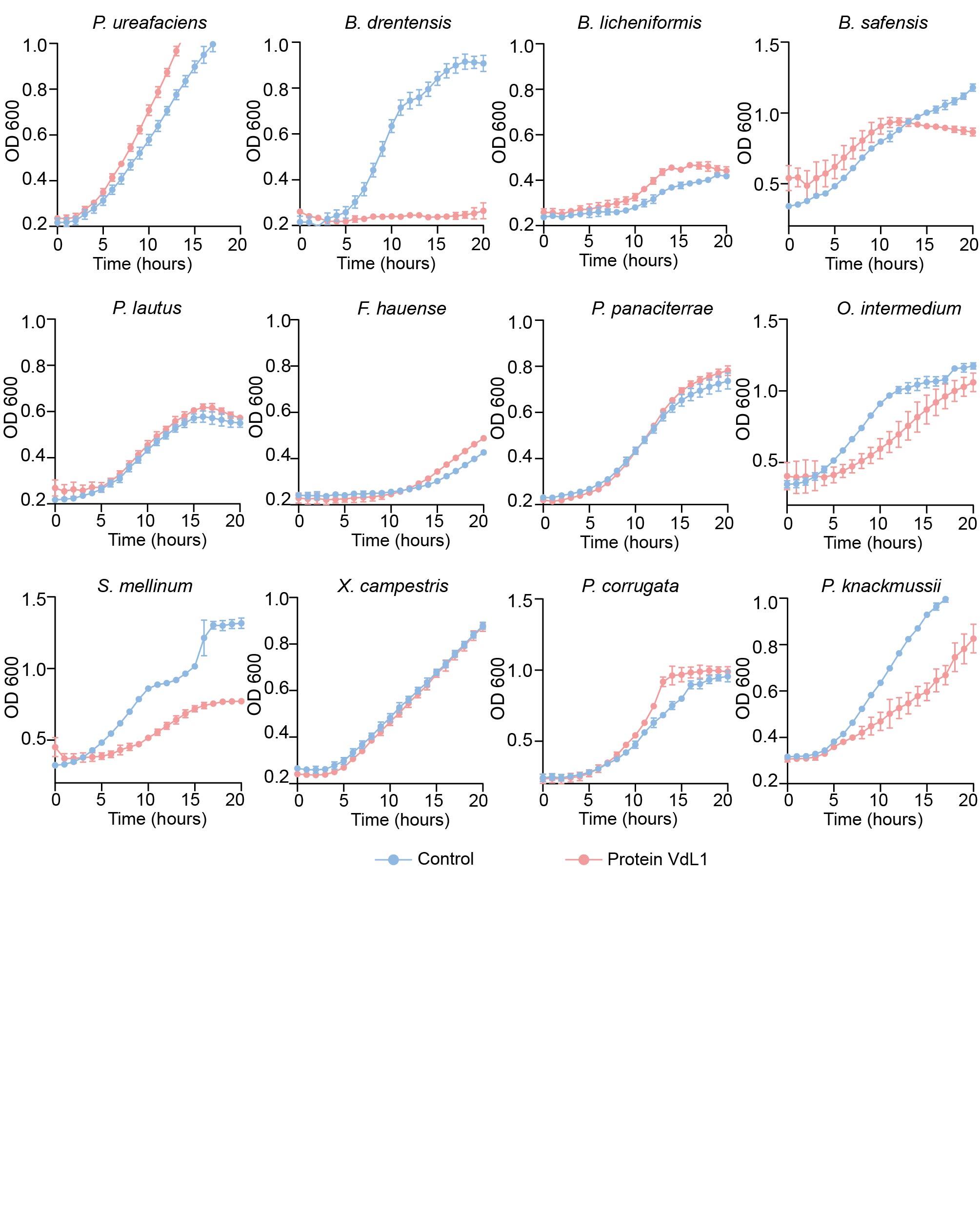

**Extended Data Fig. 8: Selective antibacterial activity of VdL1 *in vitro*.** Absorbance measurements (at wavelength 600 nm) over 20 hours of bacterial cultivation in presence and absence of 8 µM of heterologously produced VdL1. Data represent the average OD600 of three biological replicates ± SD. The assay was performed on a phylogenetically diverse set of 12 bacterial isolates, which species-level phylogeny can be seen in Fig. 5a.

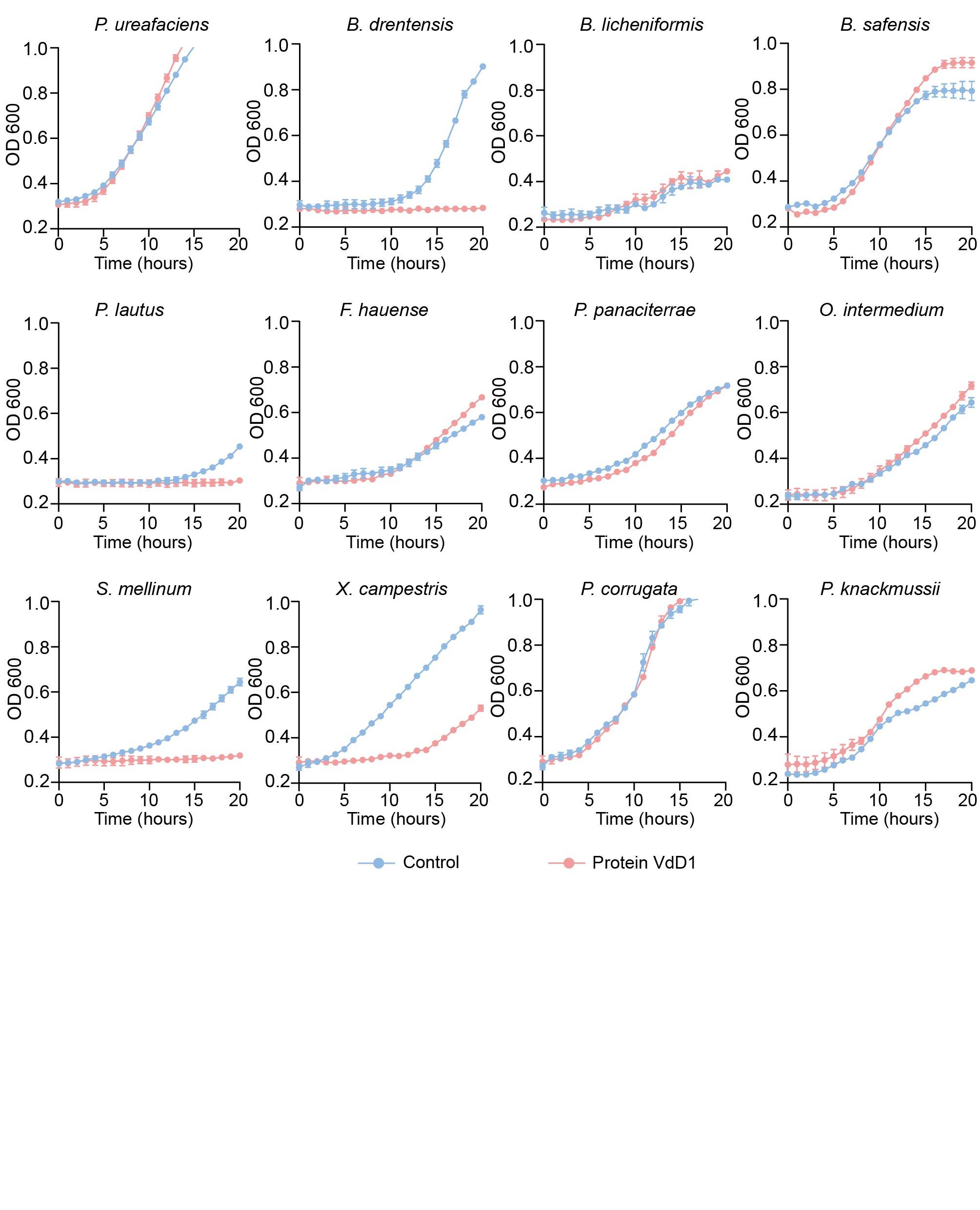

**Extended Data Fig. 9: Selective antibacterial activity of VdD1 *in vitro*.** Absorbance measurements (at wavelength 600 nm) over 20 hours of bacterial cultivation in presence and absence of 8 µM of heterologously produced VdD1. Data represent the average OD600 of three biological replicates ± SD. The assay was performed on a phylogenetically diverse set of 12 bacterial isolates, which species-level phylogeny can be seen in Fig. 5a.
